# Allele-specific analysis reveals exon- and cell-type-specific regulatory effects of Alzheimer’s disease-associated genetic variants

**DOI:** 10.1101/2021.07.26.453897

**Authors:** Liang He, Yury Loika, Alexander M. Kulminski

**Author notes:** Corresponding authors: Liang He, Alexander Kulminski.

## Abstract

Elucidating regulatory effects of Alzheimer’s disease (AD)-associated genetic variants is critical for unraveling their causal pathways and understanding the pathology. However, their cell-type-specific regulatory mechanisms in the brain remain largely unclear. Here, we conducted an analysis of allele-specific expression quantitative trait loci (aseQTLs) for 33 AD-associated variants in four brain regions and seven cell types using ~3000 bulk RNA-seq samples and >0.25 million single nuclei. We develop a flexible framework using a hierarchical Poisson mixed model unifying samples in both allelic and genotype-level expression data. We identified 24 AD-associated variants (~73%) that are allele-specific eQTLs (aseQTLs) in at least one brain region. Multiple aseQTLs are region-dependent or exon-specific, such as rs2093760 with *CR1*, rs7982 with *CLU*, and rs3865444 with *CD33*. Notably, the *APOE* ε4 variant reduces *APOE* expression across all regions, even in healthy controls. In pinpointing the cell types responsible for the observed region-level aseQTLs, we found rs2093760 as an aseQTL of *CR1* in oligodendrocytes but not in microglia. Many AD-associated variants are aseQTLs in microglia or monocytes of immune-related genes, including *HLA-DQB1, HLA-DQA2, CD33, FCER1G, MS4A6A, SPI1*, and *BIN1*, highlighting the regulatory role of AD-associated variants in the immune response. These findings provide further insights into potential causal pathways and cell types mediating the effects of the AD-associated variants.

## Introduction

Late-onset sporadic Alzheimer’s disease (AD) is the most prevalent progressive neurodegenerative disorder among all dementia cases affecting a large proportion of the elderly population (Winblad et al., 2016). Late-onset AD (LOAD) etiology is not clearly understood, posing substantive challenges of developing effective intervention and treatment procedures. With the technological breakthrough of the next-generation sequencing, large-scale genome-wide association studies (GWAS) over the past decade have hitherto detected >30 genetic loci associated with LOAD (Guerreiro et al., 2013; Harold et al., 2009; Hollingworth et al., 2011; Jansen et al., 2019; Jonsson et al., 2013; Kunkle et al., 2019; Lambert et al., 2013; Naj et al., 2011), highlighting the involvement of lipid metabolism and the immune system in the pathogenesis of LOAD. However, except for a few exonic variants in, e.g., *APOE* and *TREM2*, most of these identified signals are located in non-coding regions and thus present difficulties in elucidating their causal pathways in neuropathology and pinpointing the genes, cell types, and brain regions mediating their associations with LOAD.

A potential biological consequence of a non-coding genetic variant is its effects on regulating the expression of local genes. These variants are referred to as cis-acting expression quantitative trait loci (eQTLs). The *cis*-eQTLs have been extensively investigated in multiple tissues and regions across the brain (Katsumata et al., 2019; Park et al., 2021; Sieberts et al., 2020; The GTEx Consortium, 2020), substantially improving our understanding of the genetic role of AD-associated variants in local gene regulation. Nevertheless, these transcriptomic studies are still limited in terms of sample size, brain regions, and cell-type specificity. The statistical power is compromised by limited sample sizes and substantial inter-subject noises, including RNA degradation in the post-mortem tissues and heterogeneity of cell-type proportions (Park et al., 2021). More importantly, the brain consists of neural cell types with different morphology, and it is largely unclear which cell types mediate the effects of the eQTLs detected at the tissue level. Until now, most studies of cell-type-specific eQTLs related to AD were performed using an interaction model with deconvolved cell type proportion (Park et al., 2021; Patel et al., 2020), of which the power and accuracy could be substantially compromised. Fortunately, recent advances in high-throughput single-cell RNA-seq technologies dramatically facilitate the exploration of cell-level differential regulatory events.

To discover novel eQTLs across more brain regions and further explore their cell-type-specific regulatory effects, in this work, we focused on interrogating single nucleotide polymorphisms (SNPs) identified in GWAS of LOAD by conducting an allelic eQTL analysis with both bulk RNA-seq and single-nuclei RNA-seq (snRNA-seq) data. We investigated their *cis*-regulatory effects on the gene expression in four regions in the cerebrum using 3000+ tissue and cell-sorting bulk RNA-seq samples and >0.25 million neural cells from snRNA-seq data. Compared with a genotype-based eQTL analysis, allele-specific eQTL (aseQTL) analysis boosts the statistical power by circumventing inter-subject variation originating from confounders such as age, sex, environmental exposures, RNA degradation, and technical noises introduced in experiments. An allele-specific analysis measures the imbalance between allelic counts within each of the heterozygous subjects. This strategy is robust against such inter-subject noises, and therefore markedly improves the statistical power and the accuracy of the estimated eQTL effects.

A handful of statistical models have been proposed to leverage allelic imbalance or detect aseQTLs in bulk RNA-seq data (Degner et al., 2009; van de Geijn et al., 2015; Gutierrez-Arcelus et al., 2020; Knowles et al., 2017; Kumasaka et al., 2016; León-Novelo et al., 2014; Mohammadi et al., 2017). We classify these methods into two major categories. The methods in the first category model the allelic imbalance within each of the heterozygous samples using, e.g., a binomial model (Degner et al., 2009). More recent methods, such as a beta-binomial model, a Poisson-Gamma model (León-Novelo et al., 2014), a logistic mixed-effects model (Gutierrez-Arcelus et al., 2020), and a binomial generalized linear mixed model (Knowles et al., 2017), further account for overdispersion arising from biological and technical variations.

Despite being flexible as they can be fitted using existing standard statistical tools, these models take advantage of only heterozygous samples. The methods in the second category (van de Geijn et al., 2015; Kumasaka et al., 2016) build a dedicated joint likelihood to incorporate the contributions from both allele-specific and genotype-based information. Despite being able to include homozygous samples to improve the statistical power, the second category is less flexible for, e.g., incorporating interaction or non-linear terms.

Here, we propose a flexible and statistically powerful method based on a hierarchical Poisson mixed model (HPMM) for detecting aseQTLs. The HPMM does not only incorporate both allelic and subject-level expression but also prioritizes the heterozygous samples to boost the statistical power. Furthermore, it is easy to implement using existing statistical tools like the *lme4* R package. This is achieved by introducing hierarchical random effects and a variable coding strategy for genotypes so that allelic counts of a heterozygous sample are remodeled as paired samples. In the snRNA-seq data, unique molecular identifiers (UMIs) are often adopted to mitigate a source of bias introduced in the library amplification step during the sample preparation (Waszak et al., 2014). Such single-cell data, albeit a more accurate quantification of mRNA molecules, introduce another layer in quantifying the allele-specific expression (ASE). We then further extended this framework of HPMM to accommodate droplet-based snRNA-seq data using UMIs.

Using the HPMM, we performed a comprehensive aseQTL analysis to investigate the regulatory effects of 33 AD-associated SNPs on the expression of local genes. The early stage of LOAD involves the hippocampus and entorhinal cortex, and the degeneration spreads to other brain regions in a later stage, suggesting that the heterogeneity across brain regions may partly explain the pathogenesis of LOAD. In this study, we examined and compared the effects across the prefrontal cortex (PFC), temporal cortex (TC), posterior cingulate cortex (PCC), and head of caudate nucleus (HCN) by using bulk RNA-seq data. We further explored the cell-type-specific eQTLs and aseQTLs in five major neural cell types, including excitatory and inhibitory neurons, astrocytes, oligodendrocytes, and oligodendrocyte progenitor cells (OPCs), using snRNA-seq data comprising 240,000+ cells from the PFC. Because many AD-associated SNPs are located in or in the proximity of genes specifically expressed in microglia or monocytes (e.g., *TREM2* and *CD33*), we further evaluated the regulatory effects in these two cell types using both cell-sorting bulk RNA-seq and snRNA-seq data.

## Results

### Many AD-associated loci are aseQTLs in multiple regions in the cerebrum

We specifically focused on 33 independent (except for the *APOE* ε4 and *APOE* ε2 variants encoded by the minor alleles of rs429358 and rs7412, respectively) AD-associated common SNPs (defined as a minor allele frequency (MAF)>5%) in the European population reported or replicated in large-scale GWAS and meta-analyses (Guerreiro et al., 2013; Harold et al., 2009; Hollingworth et al., 2011; Jansen et al., 2019; Jonsson et al., 2013; Kunkle et al., 2019; Lambert et al., 2013; Naj et al., 2011) (Fig. 1a). For each of these SNPs, we examined its association with the ASE of proximal genes whose transcription starting site (TSS) is located within a window of ±500k base pairs (bp) across the four brain regions (Fig. 1b) and seven cell types (Fig. 1c, 1d). We measured SNP-level ASE through exonic variants in these local genes in bulk RNA-seq and snRNA-seq data. More specifically, we first quantified the ASE based on reads overlapping a heterozygous exonic SNP in the RNA-seq data. For the snRNA-seq data, mRNA molecules with a UMI were instead used for the quantification (See Methods). This exonic SNP is not necessarily in high linkage disequilibrium (LD) with the AD-associated SNP, but its haplotype information is available or can be estimated from, e.g., a phasing procedure during imputation. Using this haplotype information, we then employed this exonic SNP as a proxy for the gene to quantify the ASE associated with each allele of the AD-associated SNP. This SNP-level ASE can only be measured for samples with double-heterozygous genotypes at both the GWAS locus and the exonic locus. Our proposed HPMM and coding strategy (See Methods, Fig. S4) build a unified framework to further allow for incorporating subjects that are not double-heterozygous, which improves the statistical power, particularly for situations where the number of double-heterozygous subjects is low. We used WASP (van de Geijn et al., 2015) to remove potential mapping bias and included only high-quality imputed genotype data to minimize haplotype phasing errors (See Methods). We excluded from the analysis exonic loci overlapping few RNA-seq reads (a mean raw count <2) in that tissue or cell type because of their low expression and thus the lack of statistical power.

**Figure 1.**
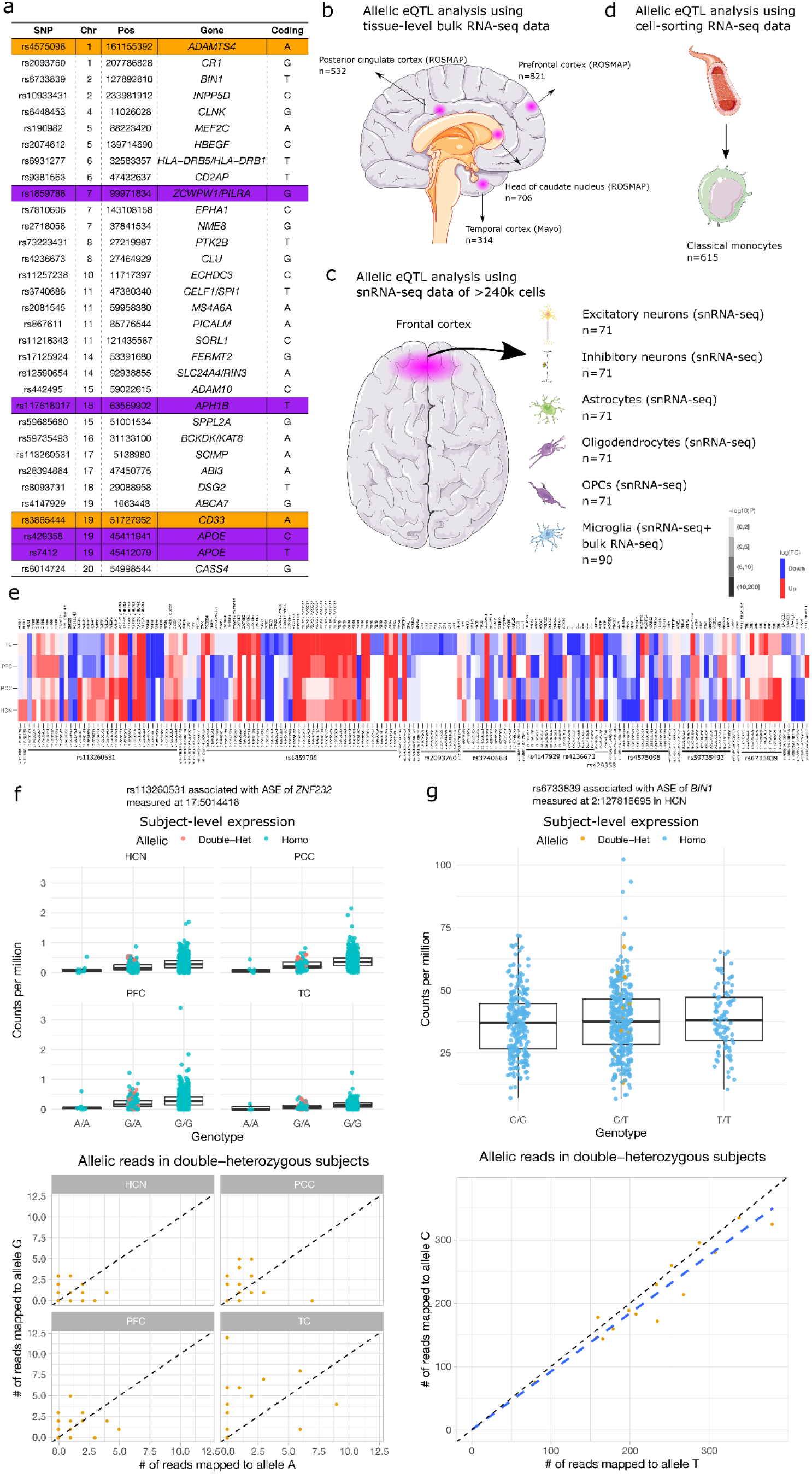
Allele-specific eQTL analysis of AD-associated loci using bulk RNA-seq and snRNA-seq data. (a) A list of AD-associated SNPs investigated in this study. Pos: the genomic position of the SNP in the hg19 human reference genome. Gene: the annotated nearest gene. Coding: the effect allele (coded as 1) for the summary statistics reported in our analysis. The UTR and missense variants are highlighted in orange and purple, respectively. The regions and cell types investigated in the aseQTL analysis (with their sample sizes) include: (b) The aseQTL analysis in four brain regions using bulk tissue-level RNA-seq data, (c) The aseQTL analysis in six neural cell types using snRNA-seq and cell-sorting RNA-seq data, and (d) The aseQTL analysis in monocytes using cell-sorting RNA-seq data. (e) Significant aseQTLs with their associated genes identified in PFC, PCC, HCN, and TC. The genomic position of the exonic variant used for measuring the ASE is given after the ID of the AD-associated SNP. The effect of up- or down-regulation is corresponding to the effect allele in (a). (f) An example of the power advantage of incorporating subjects that are not double-heterozygous. The boxplots summarize the subject-level expression, and the ASE of the double-heterozygous samples (yellow points) are shown in the scatter plot below. (g) An example of combining evidence from both ASE and genotype-level expression to improve the power for detecting the association between the AD-associated SNP rs6733839 and the ASE of *BIN1*. The dashed line is the smooth curve fitted using linear regression.

To compare the regulatory effects across different areas in the cerebrum, we performed an aseQTL analysis using four bulk RNA-seq data sets comprising ~2500 samples in the PFC, TC, PCC, and HCN from the ROSMAP (Bennett et al., 2012a, 2012b) and the MayoRNASeq Project (Allen et al., 2016) (Fig. 1b). The PCC and HCN have not been investigated in the previous studies of *cis*-eQTLs in the brain (Sieberts et al., 2020; The GTEx Consortium, 2020). We detected a total of 367 significant (false discovery rate (FDR) p<0.05) associations (a complete list of the summary statistics is provided in Table S1). These associations involve 24 (72.7%) independent AD GWAS loci in 173 haplotypes, showing differential ASE of 87 genes in at least one of the brain regions (Fig. 1e). Most of these associations showed consistent effects across the four brain regions (Fig. 1e), while some exhibited a region-specific pattern. For example, as measured by multiple coding variants in *RABEP1*, rs113260531-A was associated with reduced expression in the PFC and TC, but with elevated expression in the other regions (Fig. 1e). rs7982, in almost complete LD with the AD GWAS locus rs4236673, was significantly (p=1E-50) associated with the ASE of *CLU* only in TC, but not even nominally significant in the other regions (Fig. 1e, Table S1). In addition, rs2093760 and rs442495 were strong aseQTLs of *CR1* and *ADAM10* in TC, respectively, while the expression of both genes was hardly detectable in the other regions. These observations suggest that the regulatory effects of the AD-associated SNPs on gene expression can be region-dependent.

The advantage of the proposed HPMM to combine both allelic and genotype-level expression becomes evident in some of the findings. For example, the significance of the association between rs113260531 and *ZNF232* came primarily from the genotype-based association as the evidence for the allelic imbalance of *ZNF232* was equivocal due to the small number of double-heterozygous samples (Fig. 1f). This demonstrates the advantage of incorporating subjects that are not double-heterozygous, particularly when one of the SNPs has a low MAF. On the other hand, the significance of the association between rs6733839 and *BIN1* was achieved by combining the weak evidence from both genotype-level and allelic expression (Fig. 1g).

### SNP-level aseQTL analysis reveals exon-specific associations in *CD33* and *APOE*

The SNP-level ASE facilitates the exploration of exon-specific aseQTLs compared to haplotype-level ASE, which aggregates reads across all heterozygous loci within a gene. Opposite effects or weaker significance observed at exonic SNPs in different exons of a gene might suggest that the identified aseQTLs are splicing variants or associated with differential transcript expression. It might also be due to lower expression at the exonic variant or a smaller number of double-heterozygous subjects. Among the significant associations, we observed strong exon-specific effects at two exonic loci (chr19:51728477 and chr19: 51728641) in the second exon of *CD33* and much weaker effects in the other exons (Table S1). Because *CD33* is predominantly expressed in microglia among the neural cells (Mathys et al., 2019; Zhang et al., 2016), this region-specific result supports the previous finding of rs3865444 or its haplotype as a splicing variant of exon 2 of *CD33* in microglia (Malik et al., 2013).

Additionally, we observed exon-dependent associations in, e.g., *APOE, CLU*, and *PILRA* (Fig. 1e). These associations, however, result from the fact that the AD GWAS SNPs are in high LD with multiple exonic variants that showed opposite effects in different transcripts (Table S1). For example, we observed that the *APOE* ε4 allele (rs429358-C), the leading genetic risk factor, was associated with reduced expression of exon 4 of *APOE* (Fig. 2b), but with elevated expression of exon 2 measured at rs440446 (Fig. 2a). These effects were consistent across all the tissues in both homozygous and double-heterozygous subjects (Fig. 2a, 2b). A further investigation of the differential effects detected in these exons reveals that the *APOE* ε4 variant is in mild LD with rs440446, which is located in another transcript (ENST00000434152) of *APOE* (Fig. 2c). The expression of ENST00000434152 is much lower than that of the major transcript (ENST00000252486) (Fig. 2a, 2b), probably due to its premature termination (Fig. 2c), and ENST00000434152 does not lead to a truncated ApoE protein (Lee et al., 2020). As ENST00000252486 accounts for the vast majority of the *APOE* expression, the imbalanced ASE indicates that the haplotype containing the *APOE* ε4 allele has a repressive effect on the ApoE E4 isoform. We then attempted to elucidate whether the association between the *APOE* ε4 allele and the decreased *APOE* expression was mediated by a higher frequency of AD patients in the *APOE* ε4 carriers. We carried out an allelic analysis using the healthy controls alone in PFC, PCC, and HCN (the cohort in TC has only a small number of control subjects and thus was not investigated). We found that the significant associations were preserved in the control subjects, suggesting that the regulatory effects of the *APOE* ε4 allele were not mediated through the diagnosis of AD.

**Figure 2.**
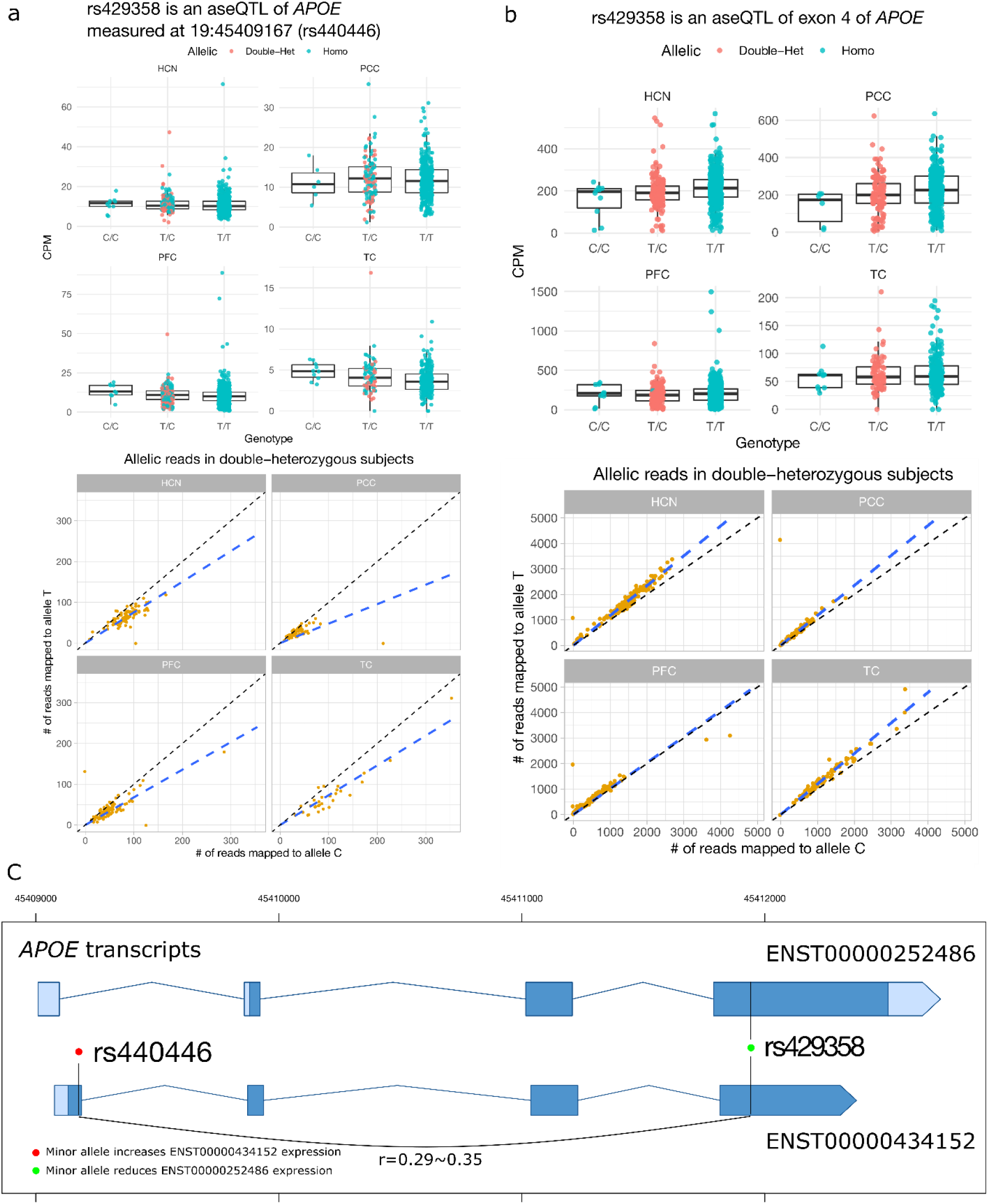
The *APOE* ε4 variant rs429358-C shows exon-dependent association with the ASE of *APOE*. (a) rs429358-C is associated with increased expression of exon 2 measured at the exonic variant rs440446 in the four brain regions. (b) rs429358-C is associated with increased expression of exon 4 measured at itself in the four brain regions. In both (a) and (b), the evidence from the subject-level expression and allelic expression is consistent. (c) The exon-dependent associations are due to the LD between rs429358 and rs440446, which had opposite effects on different *APOE* transcripts.

### Interpretation of the allelic associations by classifying aseQTLs

As shown in the example of *APOE*, the identified aseQTLs can be mediated by high LD between the AD-associated SNP and the exonic SNP, which might be the driving variant. To further explore the biological interpretation, we, therefore, divided these significant associations into three categories based on the LD between the SNPs and whether the exonic SNP per se was associated with the ASE (Fig. 3a). This strategy attempted to evaluate whether the detected allelic imbalance was driven by the AD-associated SNP or the exonic proxy SNP per se. The first category includes the situation in which the p-value of the exonic SNP is less significant than that of the AD-associated SNP for the association with the ASE (Fig. 3a, Top). We found 53 associations in this category involving the ASE of 19 genes (Table S1, Fig. 3b). The associations in this scenario are likely not mediated by the exonic SNP, and the allelic imbalance results from either the regulatory effects of the GWAS SNP or the effect of another exonic variant that is in high LD with the GWAS SNP. A notable example is those associations between rs2093760 and *CR1* in TC. The AD-associated SNP rs2093760 was significantly associated with the ASE of multiple exons of *CR1* while all exonic variants were not or were weakly associated with the ASE except for the missense variant rs2296160, which was in nearly complete LD (r=0.995) with rs2093760 and very strongly (p=8.95E-16) associated with the ASE (Table S1, Fig. 3c). This suggests that the associations between rs2093760 and the ASE of *CR1* are likely mediated by the missense variant rs2296160. This situation is described in the second category, in which the p-value of the GWAS SNP is less significant than that of the exonic SNP but the two SNPs are in high LD (defined by |r|>0.8) (Fig. 3a, Middle). We found 17 other associations in this category, including rs4236673 with *CLU*, rs4575098 with *ADAMTS4*, rs4575098 with *B4GALT3*, rs59735493 with *PRSS36*, rs3865444 with *CD33*, and rs442495 with *ADAM10* (Table S1). These AD GWAS SNPs were in almost complete LD with the exonic SNPs used for measuring the ASE. Hence, the effect can be driven by the disease-associated SNP, or an exonic SNP in high LD with it, or a haplotype harboring alleles of these two SNPs. It is not straightforward to distinguish which is the causal variant without combining additional annotation information.

**Figure 3.**
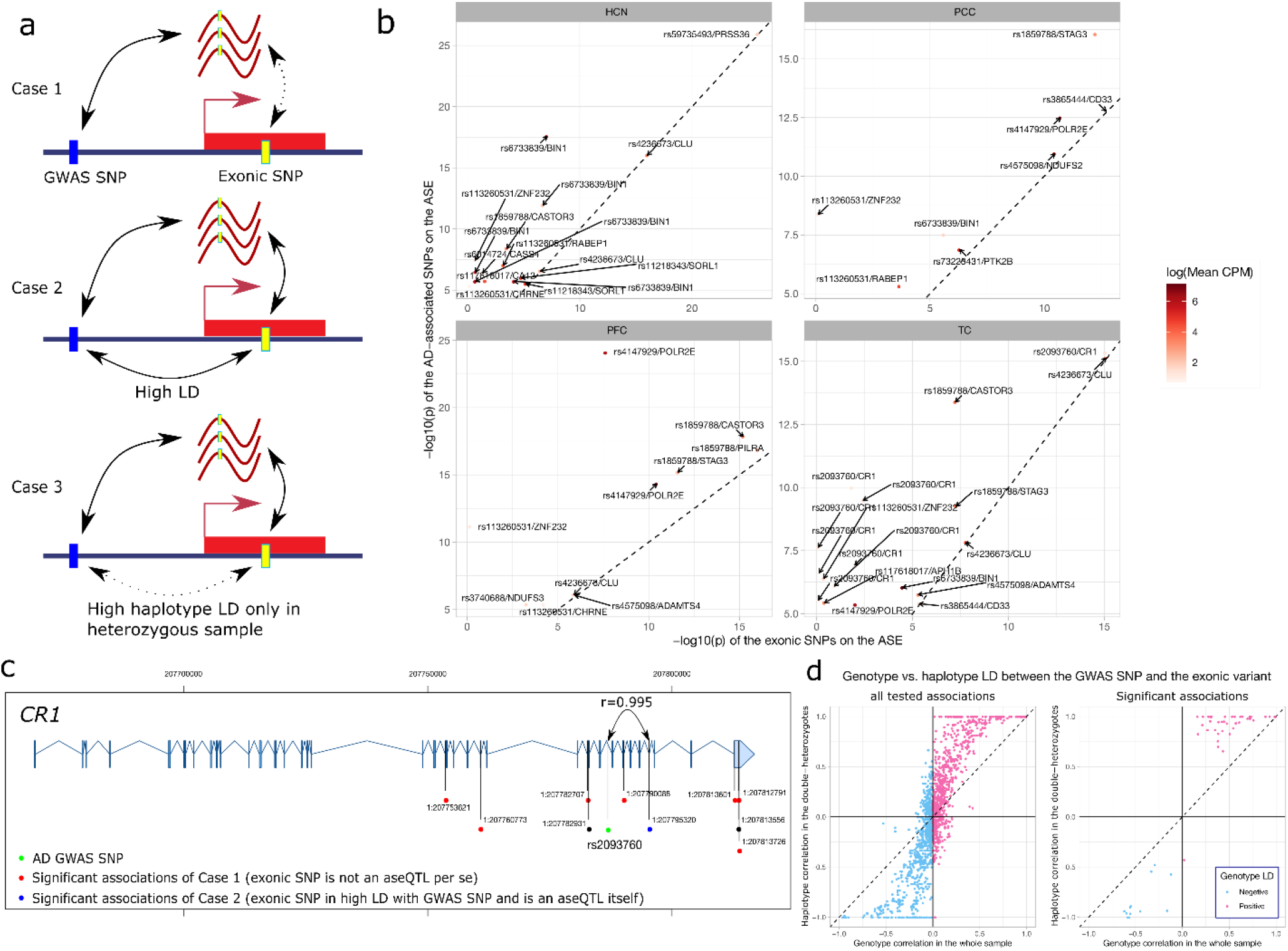
Interpretation of the identified aseQTLs in PFC, PCC, HCN, and TC. (a) We classified the associations into three scenarios based on the LD between the AD-associated and the exonic SNPs, and whether the GWAS SNP is more significantly associated with the ASE. (b) Comparison of the log(p-value) of the association with the ASE between the AD GWAS SNP (y-axis) and the exonic variant (x-axis) for the significant aseQTLs belonging to the first category. The color of the points indicates the expression level measured at the exonic variant. The labels indicate the AD GWAS SNP and its associated gene. For better visualization, we omit one significant association between rs4236673 and *CLU* in TC, which has a p-value<1E-50. (c) Exon-dependent aseQTLs identified in *CD33*. Green: the AD-associated SNP. Blue: significant associations belonging to the second category. Red: significant associations belonging to the third category. Black: Non-significant associations (d) Comparison of the genotype correlation between AD-associated and the exonic SNPs in the whole sample and the haplotype correlation observed in double-heterozygous samples.

We classify the remaining significant aseQTLs as the third category, mainly including the situation in which the disease-associated SNP was not in strong LD with the exonic SNP, and the association between the ASE and the exonic SNP was much more significant than that of the AD-associated SNP (Fig. 3a, Bottom). In this category, the allelic imbalance was probably driven by the exonic SNP, and the observed association between the ASE and the AD-associated SNP often resulted from a large haplotype LD between the two SNPs in the double-heterozygous subjects. As we showed in Fig. 3d, although the two SNPs are in very mild LD, their conditional correlation in the double-heterozygous samples can be much larger, particularly in the significant associations. This suggests that this inflated haplotype correlation among the double-heterozygous subjects plays a more prominent role as a confounder for the observed association between the ASE and the disease-associated SNP. Moreover, we observed that the difference between the haplotype correlation in the double-heterozygous subjects and the genotype correlation in the whole sample could be even more pronounced when the SNPs have a low MAF (Fig. S5) or the sample size is small (See Methods for more details). Therefore, LD has a prominent influence on detecting SNP-level aseQTLs, and it should be cautious about interpreting the associations in this category.

To further assess these significant associations, we then compared the identified aseQTLs with summary statistics of genotype-level eQTLs from a meta-analysis using the same cohorts in PFC and TC (Sieberts et al., 2020), a previous eQTL analysis in ROSMAP (Ng et al., 2017), and GTEx (v8) (The GTEx Consortium, 2020) in PFC, hippocampus, and anterior cingulate cortex (ACC). In the first category, the eQTL associations with *NDUFS2* and *APH1B* are identified in PFC in (Sieberts et al., 2020) or GTEx, and the associations with *POLR2E, ZNF232, PILRA, STAG3*, and *PRSS36* are reported in multiple regions. The association between rs2093760 and *CR1* was identified only in TC in (Sieberts et al., 2020) and the hippocampus in GTEx, corroborating our finding of its region-specific effects. In the second category, we found that *ADAM10* and *CR1* are also identified in (Sieberts et al., 2020). However, our detected aseQTLs associated with *CLU*, *BIN1, CD33, PTK2B, B4GALT3, CASTOR3*, and *ADAMTS4* in the brain are not reported in these studies. Because exonic SNPs disrupting the exon-level or transcript-level expression might not be detected from the analysis of the gene-level expression, we then compared summary statistics of splicing QTLs (sQTLs) in the three regions in GTEx and ROSMAP (Raj et al., 2018). Indeed, we found that the exonic SNPs used for measuring the ASE with *CLU*, *PTK2B, CASTOR3, BIN1*, and *CASTOR3* are in high LD with the sQTLs in the PFC reported in (Raj et al., 2018). Notably, for *CLU*, our aseQTL analysis captured two sQTLs, rs9331888, reported in (Raj et al., 2018; Szymanski et al., 2011) and rs7982, reported being associated with intron retention in exon 5 (Han et al., 2020). The AD GWAS SNP rs4236673 is in almost complete LD with rs7982, and in mild LD with rs9331888 (Table S1). The failure to detect these aseQTLs in the previous eQTL studies is probably due to its exon-specific effect, which is difficult to be captured in the gene-level analysis.

### No evidence of interaction between aseQTLs and age, sex, or AD

Because age is one of the major risk factors of LOAD and has been reported to regulate the expression of microglial genes related to cytoskeleton, immune response, and cell adhesion (Galatro et al., 2017), we further investigated whether the effects of the detected aseQTLs were different across age groups in the four brain regions. We transformed age into a dichotomized variable (age at death ≤85 or >85). We carried out an interaction analysis by adding the dichotomized age variable and its interaction term with the genotype to the HPMM. We restricted our interaction analysis to the 367 significant associations identified in the region-specific aseQTL analysis. We tested the null hypotheses of no interaction effects on the ASE between the genotype and age; that is, the effect of the aseQTL on the ASE is homogeneous across the interval of age. We found no significant interaction effects of age after adjusting for multiple testing (FDR p<0.05), suggesting that age generally does not affect the aseQTL effects in the elderly population (Table S2). In addition, we found no interaction effects of sex or AD diagnosis with genotypes for the detected aseQTLs.

### Cell-type-specific analysis reveals that rs2093760 is an eQTL of *CR1* in oligodendrocytes

Given the evidence that many AD GWAS SNPs were aseQTLs in the cortex, we next attempted to pinpoint the neural cell types mediating such associations. We collected snRNA-seq data in the frontal cortex comprising >240,000 cells from 71 subjects from two cohorts in ROSMAP (Habib et al., 2017; Mathys et al., 2019) (Fig. 1c). We classified these cells into each of the six major neural cell types, including excitatory neurons, inhibitory neurons, astrocytes, microglia, oligodendrocytes, and oligodendrocyte progenitor cells (OPCs) based on the cell type annotation in (Habib et al., 2017; Mathys et al., 2019). Unlike the bulk RNA-seq data, UMIs are utilized in these snRNA-seq data sets to mitigate the amplification bias. Since multiple reads tagging with the same UMI originate from the same mRNA molecule, the UMI-based snRNA-seq data introduce another layer of complication in the allele-specific analysis. We, therefore, called the ASE at the UMI level by collapsing all reads sharing a similar UMI barcode (see the Methods section for more details). We carried out a cell-type-specific aseQTL analysis for the AD-associated SNPs in each of the cell types using the UMI-level ASE of exonic variants in local genes whose TSS is located within a window of ±500k bp surrounding the AD-associated SNPs.

We tested 499 associations between 28 AD-associated SNPs and 118 local genes showing at least moderate expression (mean count>2), among which 327 tests (65.5%) were in the neurons (Fig. 4a). Only four of these tests were in microglia because of the tiny proportion of microglia in the entire library of the snRNA-seq data. Compared to the bulk RNA-seq data, we observed substantially attenuated expression in the 5’-end, which is expected because of the bias of read coverage towards the 3’-end in the 3’-sequencing snRNA-seq data. Therefore, most exonic variants investigated in this cell-type-specific analysis were located in the 3’ untranslated region (UTR) (Fig. 4b) because the reads in the other genomic regions are often not sufficiently abundant for measuring the ASE. We identified 11 significant associations (FDR p<0.05) between two AD-associated SNPs and the ASE of three genes, including *PILRB, TRIM4*, and *MTCH2* (Table S3). All these associations were also detected in the region-specific analysis (Fig. 1e). The results from the snRNA-seq data provide more details about the underlying cell types responsible for the association. More specifically, the AD GWAS SNP rs1859788 was associated with the allelic imbalance of *PILRB* in excitatory and inhibitory neurons, astrocytes, and oligodendrocytes. The trend of the genotype-level expression in the homozygous subjects is concordant with the imbalance of the ASE (Fig. 4c, 4d). These associations belong to the third category described above. rs1859788 was in complete LD with the two exonic SNPs only in the double-heterozygous subjects, while the correlation was low in the whole sample (r=~0.2), and both exonic SNPs were much more strongly associated with the allelic imbalance of *PILRB* (Table S3). Therefore, these associations are likely mediated by the effects of the two exonic SNPs and not driven by the regulatory effects of rs1859788. Similar situations were observed for the other two associations between rs1859788 and *TRIM4*, and rs3740688 and *MTCH2* in the excitatory neurons (Table S3, Fig. S3). We further investigated these identified associations by performing a general genotype-level eQTL analysis for rs1859788 and rs3740688. We found that these associations were not significant in the general eQTL analysis (Table S4), suggesting that the observed associations in the ASE analysis are likely mediated by the effects of the exonic SNPs.

**Figure 4.**
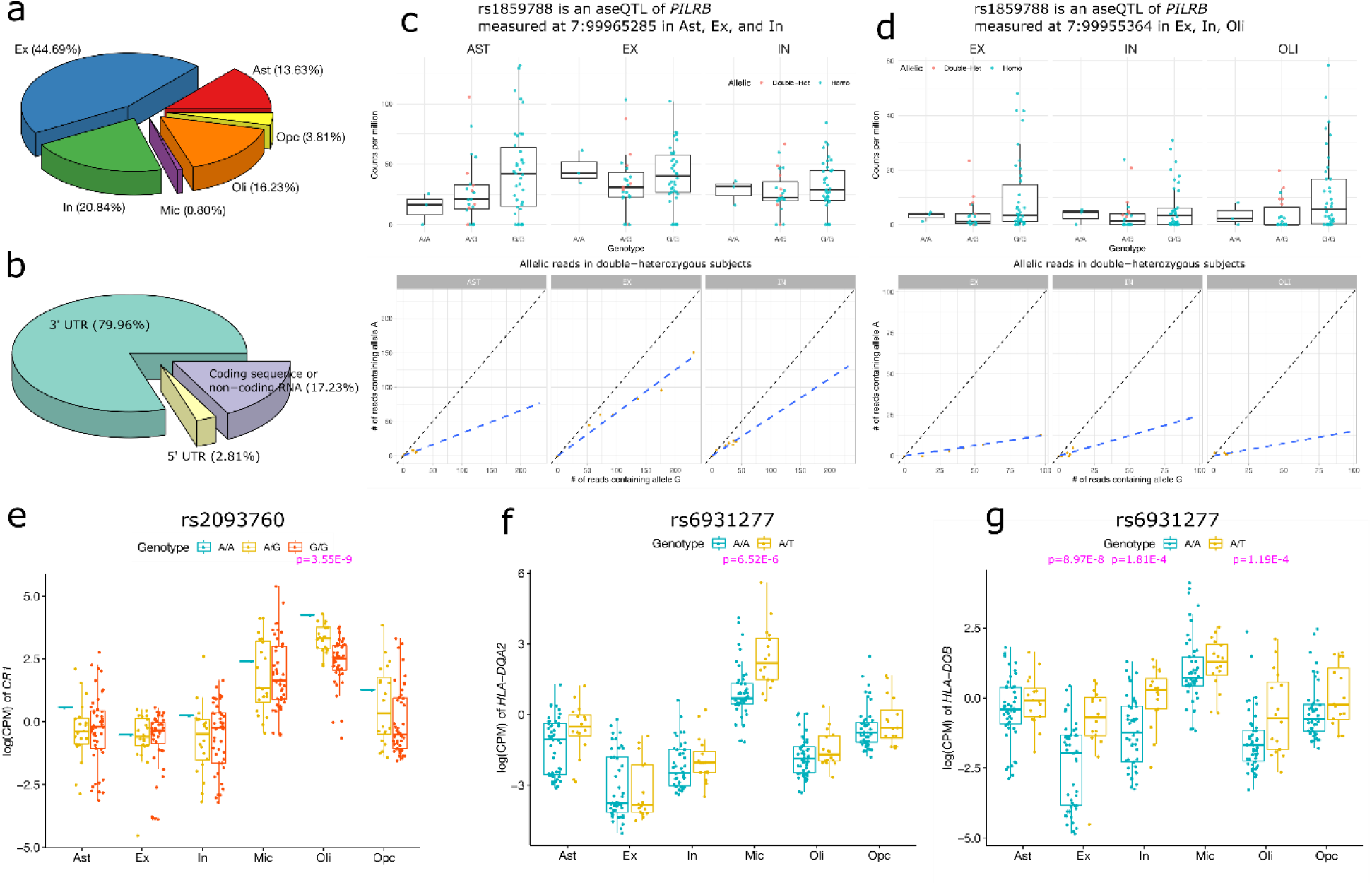
Cell-type-specific aseQTL analysis in frontal cortex using snRNA-seq data. (a) Percentage of the associations examined in the six neural cell types in the analysis of the cell-type-specific aseQTLs. (b) Percentage of the exonic SNPs located in 3’ UTR, 5’ UTR, and coding regions in the analysis of the cell-type-specific aseQTLs. (c) and (d): rs1859788 is a significant aseQTL of *PILRB* measured at two exonic loci in multiple neural cell types. The boxplots summarize the subject-level expression, and the ASE of the double-heterozygous samples (yellow points) are shown in the scatter plot below. The dashed line is the smooth curve fitted using linear regression. (e) – (g): significant genotype-based *cis*-eQTLs among the AD-associated SNPs identified using a pseudo-bulk sample by aggregating cells in the snRNA-seq data.

A drawback of the allelic analysis in the snRNA-seq data is the low expression called at exonic variants that are not close to 3’-end, which potentially misses many signals and leads to a small number of identified associations. To compare the results more thoroughly between the allele-specific and the genotype-level eQTL analyses, we further performed a genotype-level *cis*-eQTL analysis of the AD GWAS loci for cis-genes within a 500k bp window in each of the six cell types. We identified three significant associations (FDR p<0.05) (Table S4), including an association between rs2093760 and *CR1* in the oligodendrocytes (Fig. 4e). We found that *CR1* was abundantly expressed only in oligodendrocytes and microglia (Fig. 4e), but no association was detected in microglia. This suggests that the association between rs2093760 and the expression of *CR1* observed in the above tissue-level eQTL analysis is likely attributed to its effect in oligodendrocytes. We also found associations between rs6931277 and *HLA-DOB* and *HLA-DQA2* in the excitatory neurons and microglia, respectively (Fig. 4f, 4g). The association between rs6931277 and *HLA-DOB* was also nominally (p<0.05) significant in the inhibitory neurons and oligodendrocytes.

### Multiple AD-associated SNPs are aseQTLs in microglia

Many AD-associated loci are located in genes that are exclusively or abundantly expressed in microglia. Unfortunately, microglia accounted for only a small fraction of the total cell population in the two snRNA-seq data sets in the frontal cortex. To increase the statistical power to detect aseQTLs in microglia, in addition to the 71 snRNA-seq samples in the frontal cortex, we further added seven samples from a microglia-specific snRNA-seq data set (Olah et al., 2020), and ten samples from a cell-sorting bulk RNA-seq data set (Olah et al., 2018). We tested 183 associations between the imbalance of ASE that had the average count >2 and haplotypes that had at least three double-heterozygous samples. We identified seven significant associations (FDR p<0.05) (Table S5), involving six genes (*BIN1*, *ITGAM, FCER1G, MS4A7, ATXN3*, and *ABCA7*). The associations with *BIN1, FCER1G*, and *ABCA7* were also observed in our region-specific aseQTL analysis (Fig. 1e), suggesting that some of these tissue-level associations might be driven by their effects in microglia. For example, because microglia are the primary cell type expressing *FCER1G*, its tissue-level association is likely mediated through microglia. For these three associations, the ASE and the genotype-level expression exhibited consistent trends (Fig. 5a, 5b, 5c). The other three associations (*ITGAM, MS4A7*, and *ATXN3*) were not detected in the region-specific analysis, probably due to the much lower expression of these genes at the tissue level. Most significant associations in microglia were primarily driven by strong allelic imbalance observed in the double-heterozygous samples (Fig. 5a, 5c, Fig. S2). Because of the small sample size, the allelic imbalance can be attributed to an inflated correlation between the AD-associated SNP and the exonic SNP in the double-heterozygous subjects. Further study with larger sample size is required to confirm these findings.

**Figure 5.**
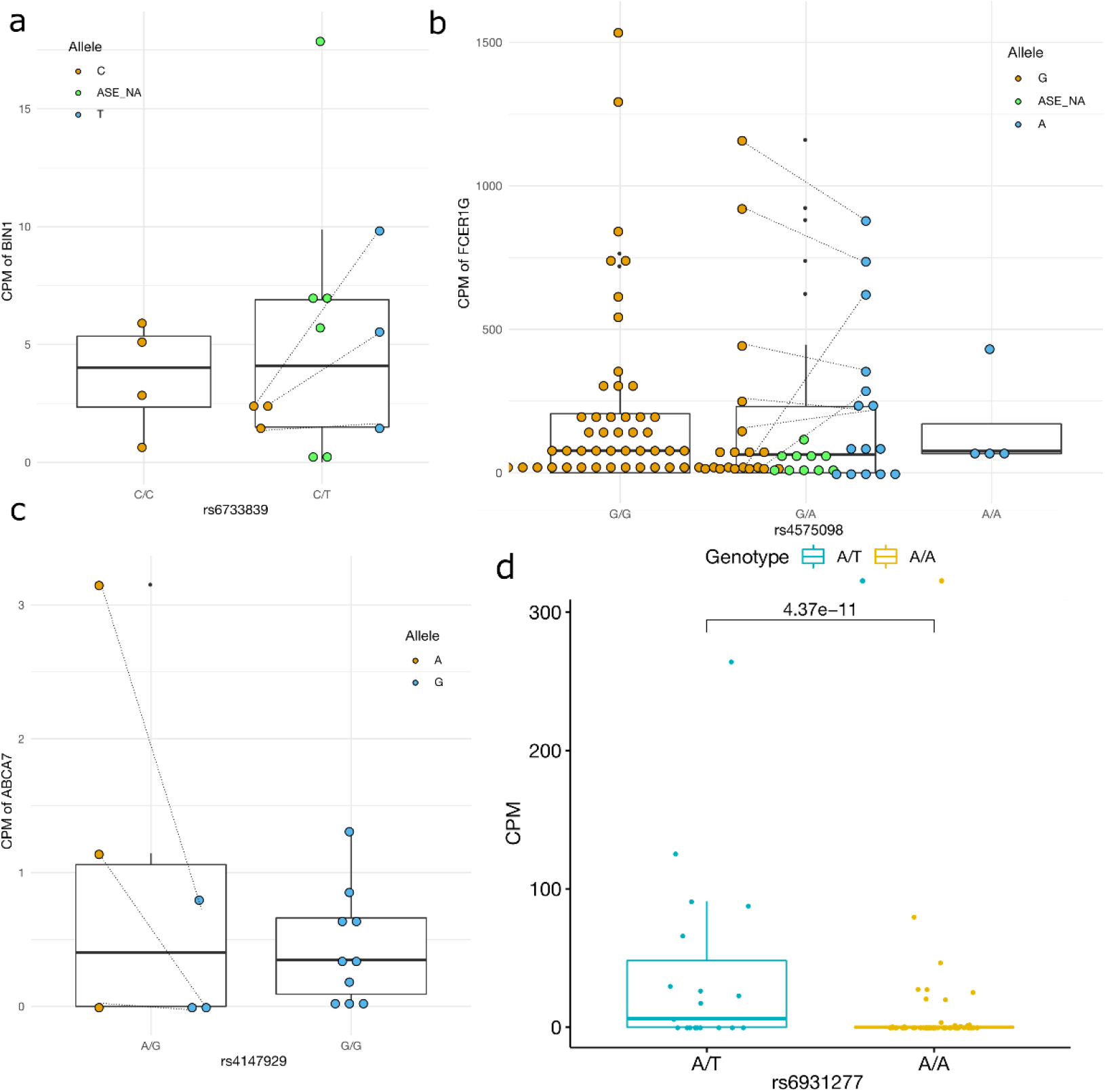
Cell-type-specific aseQTL analysis in microglia by combining snRNA-seq and cell-sorting bulk RNA-seq data sets. (a) – (c): Three significant aseQTLs identified in both microglia and the tissue-level aseQTL analysis. ASE_NA indicates those samples that ASE is not available because they are heterozygous but not double-heterozygous. The pair of the ASE of a double-heterozygous sample is connected by a dashed line. Some lines are omitted from the plot of *FCER1G* to improve visibility. The CPM of the ASE of the double-heterozygous sample was normalized by half of its total library size. (d) rs6931277 is a significant genotype-level cis-eQTL of *HLA-DQA2* in microglia.

To compare with the allele-specific analysis, we also performed a genotype-level eQTL analysis using the combined 88 samples, including the 17 microglia-specific samples and the 71 samples comprising cells annotated as microglia in the two snRNA-seq data sets in the frontal cortex. Our analysis confirmed that rs6931277 was a *cis*-eQTL of *HLA-DQA2* in microglia with a more significant p-value after including the microglia-specific samples (Fig. 5d, Table S6). In addition, among the top five genes, we observed associations between rs4575098 and the expression of *ADAMTS4* (p=7.4E-4), and between rs867611 and *PICALM* (p=9E-4) (Table S6). In contrast to the oligodendrocytes, rs2093760 was not associated with the expression of *CR1* in microglia. These associations were preserved after adjusting for the AD diagnosis (Table S6), suggesting that these cis-regulatory effects are not affected by the disease status.

### Identification of aseQTLs with immune-related genes in monocytes

Because multiple putative AD-related genes are also abundantly expressed in monocytes, which might also be involved in AD by, e.g., infiltration through the blood-brain barrier (Martin et al., 2017; Saresella et al., 2014; Young et al., 2019) or its breakdown (Montagne et al., 2015; Nation et al., 2019), we next investigated the association between the AD-associated loci and the ASE in CD14^+^CD16^-^ classical monocytes in a cohort (Olah et al., 2018) comprising >600 cell-sorting RNA-seq samples from ROSMAP (Fig. 1d). We identified 14 significant associations (FDR p<0.05) involving 6 AD-associated SNPs and 9 genes (Table S7). Most of these genes are related to the immune system, including *CD33*, *MS4A6A, SPI1, HLA-DQB1*, and *FCER1G*. In these associations, the GWAS loci were in moderate or high LD (|r|>0.5) with the exonic variants.

Consistent with the findings in the region-specific analysis, the AD GWAS SNP rs3865444 was associated with the ASE of the second exon of *CD33* measured at rs2455069 and rs12459419 in monocytes (p=4.52E-12 and 2.05E-6) (Fig. 6a), confirming the previous findings of the sQTL of exon 2 in monocytes (Raj et al., 2014). We further performed a transcript-level eQTL analysis for *CD33*, and confirms that rs3865444 was only associated with two transcripts (ENST00000421133 and ENST00000601785) (Table S8), which overlap the second exon among all seven transcripts of *CD33*.

**Figure 6.**
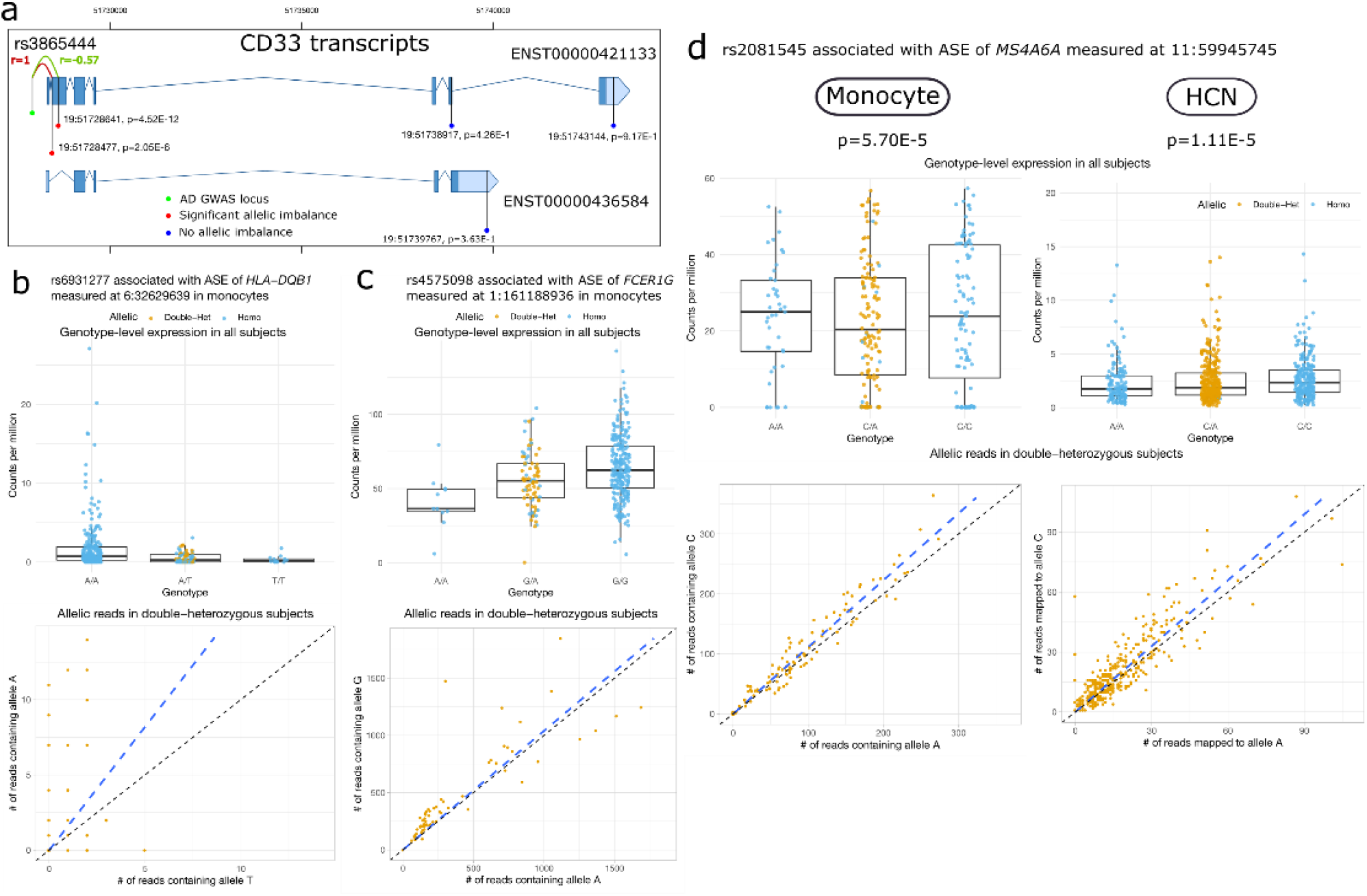
Analysis in monocytes using a cell-sorting bulk RNA-seq data set identified aseQTLs that are also observed in microglia and the brain regions. (a) Exon-specific aseQTLs of *CD33* identified in monocytes. Green: the AD-associated SNP rs3865444. Red: loci with a significant FDR p-value (FDR p<0.05). Blue: loci with a non-significant p-value (p>0.05). The red and green curves indicate complete and moderate LD between rs3865444 and the two exonic variants in exon 2 of *CD33*. P-values: the p-values of the association between rs3865444 and the ASE measured at the exonic locus. (b) rs6931277 is a significant aseQTL of *HLA-DQB1* in monocytes, which is also identified in neurons and microglia. (c) rs4575098 is a significant aseQTL of *FCER1G* in monocytes, which is also identified in microglia and multiple regions in the brain. (d) rs2081545 is a significant (FDR p<0.05) aseQTL of *MS4A6A* in monocytes, which is consistent with its p-value and effect in HCN (FDR p<0.1). The boxplots summarize the subject-level expression, and the ASE of the double-heterozygous samples (yellow points) are shown in the scatter plot. The dashed line is the smooth curve fitted using linear regression.

In addition, many associations showed consistent effects compared to those detected in microglia and the brain regions. Different from the gene *HLA-DQA2* identified in the neurons, rs6931277 was associated with both the expression and ASE of *HLA-DQB1* in monocyte (Fig. 6b). This SNP was also associated with *HLA-DQB1* and *HLA-DQA2* in microglia (Table S5) and has been reported as an sQTL of multiple Class II HLA genes in almost all tissues in GTEx (The GTEx Consortium, 2020). This suggests that rs6931277 may affect different genes in a cell-type-dependent manner. rs4575098, an AD GWAS SNP located in the UTR of *ADAMTS4*, was in moderate LD (r=0.59) with an exonic variant rs11421, located in the 5’ UTR of *FCER1G*, which was also a significant aseQTL of *FCER1G* (Fig. 6c). This association was also consistent with those detected in the region-specific analysis and in microglia in terms of the direction of the effect (Table S1, Table S5). The association between rs2081545 and the ASE of *MS4A6A* was observed in two of the exons measured at rs7232 and rs12453. The lack of significance in the other exons was probably due to their much lower expression (Table S7). The association between rs2081545 and *MS4A6A* was also observed in HCN (FDR p<0.1), and the direction of effect sizes was consistent (Table S1, Fig. 6d).

The evidence from allelic imbalance is in line with that from genotype-level differential expression in most of these associations (Fig. 6b, 6c, 6d, Fig. S1). Allelic imbalance plays a more crucial role in determining the association when the conclusion cannot be drawn from genotype-level expression alone (e.g., *MS4A6A* and *HBEGF*). In comparison, a previous eQTL analysis in classical monocytes using the DICE (~100 samples) (Schmiedel et al., 2018) identifies associations with four SNPs, one of which between rs6931277 and *HLA-DQB1* was also observed in this analysis. The other associations were not investigated in this study because these genes were not within our defined window for a *cis*-gene.

## Discussion

In this study, we investigated the regulatory effects of AD GWAS loci on the allelic expression of local genes in four regions in the brain and in six major neural cell types and monocytes. Our results, showing that most AD-associated loci are eQTLs in the brain, replicate not only previously reported signals, and, most importantly, also discover novel aseQTLs. (Sieberts et al., 2020) show that the regulatory patterns of eQTLs are different between the cortex and the cerebellum. Our SNP-level allele-specific eQTL analysis further reveals that the regulatory effects of AD-associated SNPs can be heterogeneous across different brain regions and neural cell types, even within the cortex, and can be exon-specific. Using the snRNA-seq and cell-sorting bulk RNA-seq data, we further pinpointed the cell type that mediates the eQTL effects observed at the tissue level.

We have demonstrated that the SNP-level aseQTL analysis not only identifies eQTLs but can also be used to uncover exon-specific regulation. As an example, our aseQTL analysis detected allelic imbalance at two exonic SNPs in exon 2 but not in the other exons. These associations were observed in monocytes and multiple brain regions, including TC, PCC, and HCN, supporting previous findings showing that the AD GWAS locus rs3865444 is associated with the splicing of exon 2 of *CD33* (Malik et al., 2013; Raj et al., 2014, 2018). The almost complete LD between the promoter variant rs3865444 and the exonic variant rs12459419 makes it challenging to determine the causal variant. Another example is *CLU*, of which our aseQTL analysis captured two sQTLs in the promoter and exon 5. A recent study reveals that rs7982 is associated with intron retention in three regions in the temporal lobe, including TC, superior temporal gyrus, and parahippocampal gyrus (Han et al., 2020). Our result shows no association in PFC, PCC, or HCN, suggesting that this sQTL is region-dependent.

Interestingly, our findings reveal that the *APOE* ε4 variant is associated with reduced expression of the major transcript of *APOE* based on the evidence from both allelic imbalance and gene-level expression, corroborating previous findings of lower mRNA and protein levels of *APOE* in ε4 carriers (Bertrand et al., 1995; Gottschalk et al., 2016; Poirier, 2005; Sullivan et al., 2011). This association remains significant in the control subjects and is across all the regions. The expression of the ApoE E4 isoform is lower than that of the ApoE E3 isoform in *APOE* ε3/ε4 carriers at the tissue level. Because *APOE* expresses most abundantly in astrocytes among the neural cell types, astrocytes might be the primary cell type contributing to this effect. The higher proportion of *APOE* ε4 carriers among AD patients can lead to a decreased expression in AD patients due to its eQTL effect, explaining the reported association between the reduced *APOE* expression in astrocytes and the diagnosis of AD (Mathys et al., 2019). Compared with astrocytes, the *APOE* expression in microglia is upregulated in AD patients (Mathys et al., 2019). The upregulation of *APOE* in microglia can result from either a disrupted regulation specifically in microglia or a disease-associated subpopulation, although recent studies have not detected an AD-specific subpopulation in microglia in the cortex of AD patients (Alsema et al., 2020; Mathys et al., 2019). More studies are needed to examine this regulatory effect of the *APOE* ε4 variant at the cell level, including microglia.

Using the single-cell data, we have attempted to pinpoint the cell type that mediates genetic regulatory effects detected at the region-specific analysis. For example, the association between the haplotype containing the GWAS SNP rs2093760 and the ASE of *CR1* observed in the TC was seen in oligodendrocytes, but not in microglia. *CR1* is one of the key genes involved in the complement system, and shows the highest expression level in oligodendrocytes followed by microglia among the neural cells in our data. Genes in the complement system are abundantly expressed in microglia, and are co-expressed with *APOE* (He, 2021). Our findings suggest that oligodendrocytes might play a role in mediating the effect of rs2093760. Nevertheless, this haplotype also includes rs2296160, a nonsynonymous variant of *CR1* in almost complete LD with rs2093760. More research is needed to elucidate which functional consequence of this haplotype is the primary factor contributing to the risk of LOAD. Compared to a previous study detecting few *cis*-eQTLs in the primary microglia (Young et al., 2019), we identified multiple aseQTLs and eQTLs in microglia, such as *BIN1, FCER1G, ABCA7*, and *HLA-DQB1*, which were also detected in the region-specific analysis. These findings suggest that microglia might mediate their effects observed at the tissue level.

We also proposed an HPMM to further increase the statistical power by prioritizing the evidence from the allelic imbalance and also combining the evidence from both double-heterozygous and homozygous subjects. This method is fast, easy to implement, flexible for adjusting additional covariates and investigating interactions. Our real data analysis also demonstrated that more associations were detected after combining evidence from both subject-level expression and the ASE of the double-heterozygous subjects. The additional information from the subject-level expression is important, especially when the evidence from the ASE is equivocal due to the small sample size. Future work can be focused on further extending the HPMM to accommodate haplotype-based ASE.

It should be noted that the LD between the GWAS SNP and the exonic SNP among the double-heterozygous subjects complicates the interpretation of the findings from the SNP-level aseQTL analysis. It is easier to draw a conclusion when the exonic SNP used as the proxy for measuring the ASE is not associated with the ASE intrinsically. Otherwise, we show both theoretically and practically that a false interpretation of the regulatory function can result from an inflated conditional correlation between the exonic proxy SNP and the GWAS SNP among double-heterozygous subjects. We implement a straightforward classification strategy to tackle this issue to gain better biological insights. This problem might not be serious in a haplotype-level allele-specific analysis because the ASE was quantified by aggregating multiple exonic SNPs of the gene.

One of the limitations of the cell-type allele-specific analysis using the 3’-sequencing snRNA-seq data is that the reads are distributed non-uniformly, and its concentration is biased towards 3’ UTR or 5’ UTR depending on the experimental protocol. Consequently, many exonic SNPs within the gene body are not captured by most reads or covered by a quite low number of reads, which reduces the statistical power to detect associations in these regions. For this reason, we removed loci with a low read count. More sophisticated methods based on shrinkage might be utilized to analyze these low-count loci (Zitovsky and Love, 2020). Another limitation of this study is the small sample size in the snRNA-seq data, which might at least partly justify the lack of significant findings in the cell-type-specific analysis.

In conclusion, a large number of AD GWAS loci show strong associations with the expression of local genes or exons, which can be tissue- or cell-type-dependent. Our results pinpointed the underlying brain regions and cell types that mediate the associations for multiple AD GWAS loci, which provide valuable insights into the cellular regulatory mechanisms in the genetic architecture underlying the pathogenesis of LOAD.

## Methods

### Sample collection

We obtained 747, 572, and 882 bam files of the tissue-level bulk RNA-seq samples in PFC, PCC, and PFC, respectively, in ROSMAP from Synapse (https://www.synapse.org/#!Synapse:syn22333035). The library was generated using either a poly-A selection or ribosomal depletion method. The reads were trimmed before being aligned to the reference genome (hg19) using Bowtie. These RNA-seq samples were collected from 925 subjects with an age of death ranging from 67.4 to 108.3, among whom 602 are female, and 571 are AD patients diagnosed by the pathology of the post-mortem brain. The bam files of the 317 bulk RNA-seq samples in TC from the MayoRNAseq project (Allen et al., 2016) were downloaded from Synapse (https://www.synapse.org/#!Synapse:syn20818651). This data set included 82 AD patients, 84 progressive supranuclear palsy (PSP) patients, 71 subjects with pathologic aging, and 78 healthy controls. The reads were aligned to the GRCh38 reference genome using the SNAPR software (Magis et al., 2015).

We obtained the fastq files of the cell-sorting bulk RNA-seq data sets in monocytes and microglia in ROSMAP from Synapse (https://www.synapse.org/#!Synapse:syn11468526 and https://www.synapse.org/#!Synapse:syn22024496). The monocytes were extracted based on the surface marker CD14^+^CD16^-^ from peripheral blood mononuclear cells (PBMCs) of 615 subjects. We included 336 samples that had both the RNA-seq and genotype data, including 238 female and 139 AD patients. RNA molecules were extracted using the ribosomal depletion method, and the library was prepared using a SMART-seq2 or SMART-seq2-like protocol. The microglia were isolated from the dorsolateral PFC of 10 subjects using magnetic anti CD11b beads and then followed by fluorescence-activated cell sorting based on the CD11b^high^CD45^int^7AAD^-^ staining profile. RNA molecules with a poly-A tail were extracted, and the library was constructed using the SmartSeq-2 protocol (Picelli et al., 2014). More details about the data generation are given in (Olah et al., 2018).

We obtained the fastq files of the three snRNA-seq data sets, two in the PFC and one in microglia, in ROMSAP from Synapse (https://www.synapse.org/#!Synapse:syn16780177, https://www.synapse.org/#!Synapse:syn12514624, and https://www.synapse.org/#!Synapse:syn23446265). The two snRNA-seq data sets in the PFC included ~80,000 cells in Brodmann area (BA) 10 from 48 subjects and >160,000 cells from 24 subjects. More details about the generation of these two data sets are described in (Habib et al., 2017; Mathys et al., 2019). In the snRNA-seq data set in microglia, the tissue was extracted from the dorsolateral PFC (BA9/46) of 13 subjects. The microglia were isolated using the same procedure as in the above cell-sorting bulk RNA-seq data. More details about this data set can be found in (Olah et al., 2020). All these snRNA-seq libraries were generated using the 10x Genomics Chromium Single Cell 3’ Reagent Kit v2 protocol.

Clinical characteristics of the subjects in ROSMAP and the MayoRNAseq project, including sex, age, race, ethnicity, type of neurodegenerative disorder, and age at death, were also retrieved from Synapse.

### Quantification of ASE in Bulk RNA-seq data

In the two bulk RNA-seq data sets in microglia and monocytes that started with fastq files, we first trimmed the paired-end reads using Trim Galore! (https://www.bioinformatics.babraham.ac.uk/projects/trim_galore/). The trimmed fastq files were then aligned to the hg19 human genome assembly using the STAR aligner (Dobin et al., 2013). Samples with a mapping rate <50% were excluded from the downstream analysis. Gene-level raw counts were quantitated from the bam file for each sample using the Rsubread package (Liao et al., 2019) with the parameters “requireBothEndsMapped=TRUE, allowMultiOverlap=TRUE, countMultiMappingReads=FALSE, largestOverlap = TRUE” and the gene transfer format (GTF) build GRCh37.87. In the allele-specific analysis, mapping bias introduced during the library alignment is one source for false positives (Panousis et al., 2014).

We used WASP (van de Geijn et al., 2015) in the STAR aligner to tag and remove reads that showed the alignment bias at exonic SNPs. The exonic SNPs considered in our analysis were defined by those that were included in NCBI dbSNP Build 144 (https://bioconductor.org/packages/release/data/annotation/html/SNPlocs.Hsapiens.dbSNP144.GRCh37.html) and the Haplotype Reference Consortium (HRC) SNP reference panel, and intersect at least one exon defined in the GTF build GRCh37.87. Using the WASP-filtered bams files, the ASE at the exonic SNPs were quantified using the pileup function in the samtools (Li et al., 2009) with the parameters “min_base_quality = 10L, min_mapq=254, distinguish_strands=FALSE, max_depth=10000, include_insertions=TRUE, include_deletions=TRUE” to filter out low-quality and ambiguously mapped reads.

Since the RNA-seq data sets in PFC, PCC, HCN, and TC are available only in bam files, we re-aligned the reads using WASP to control potential mapping bias in the allele-specific analysis. More specifically, we first removed secondary alignments from the original bam files using samtools and re-paired the reads according to the fragment ID using Rsubread. We then used this re-paired bam file as the input in the STAR aligner. The quantification of ASE then followed the same steps described above.

### Quantification of ASE in snRNA-seq data

The raw fastq files from the 10x Genomics platform were processed using STARsolo (Kaminow et al., 2021) to obtain the bam files aligned to the hg19 human reference genome. We used WASP to tag and filter those reads that overlapped the exonic SNPs and showed mapping bias. In STARsolo, we used the parameters “--soloCellFilter EmptyDrops_CR --soloCBmatchWLtype 1MM_multi_Nbase_pseudocounts --soloUMIfiltering MultiGeneUMI_CR --soloUMIdedup 1MM_CR” to filter cells and collapse UMI barcodes. We confirmed that this setting generated highly consistent results with CellRanger v3. We first quantified the UMI-level ASE at each of the exonic SNPs for each cell. Given a cell barcode and an exonic locus, we obtained all UMI barcodes that had at least one read overlapping the locus. Then for each of the UMI barcodes, we used the same pileup function as described above to count the number of allelic reads that had that UMI and overlapped the locus. We counted only those reads assigned to a transcript by STARsolo. We removed problematic UMIs to which multiple reads were mapped but had different alleles at the exonic locus because this may indicate a sequencing error in one of the reads. Next, we collapsed the UMIs that had at most one mismatch and obtained allele-specific counts by counting the number of collapsed UMIs that shared the allele. This gave us the UMI-level ASE in each cell at each exonic locus of interest. Finally, we aggregated all cells annotated to the same cell type and subject to obtain the cell-type-specific subject-level UMI-level ASE, which was used in the downstream allele-specific eQTL analysis.

### Genotype data processing and imputation

The three genotype data sets in ROSMAP were downloaded from Synapse (https://www.synapse.org/#!Synapse:syn17008936), including a whole-genome sequencing (WGS) data set of 1196 subjects in the VCF format and two SNP array data sets of 382 (Illumina HumanOmniExpress) and 1709 (Affymetrix GeneChip 6.0) subjects in the plink format. If a subject had both the WGS and SNP array data, we always used the WGS data for more accuracy. Before the imputation, we checked strands, alleles, and positions and removed ambiguous SNPs using the tool HRC-1000G-check-bim.pl with default settings and the reference panel of the EUR population in the 1000 Genomes project (https://www.well.ox.ac.uk/~wrayner/tools/). We phased and imputed the WGS data set using the Michigan Imputation Server (Das et al., 2016) with the HRC reference panel (Version r1.1 2016). Because one of the SNP array data sets using the Affymetrix GeneChip 6.0 has many missing data and low-quality genotypes, after removing the subjects and SNPs with a missing rate >5%, 1019 subjects remained. To achieve better imputation quality for the SNP array data, we used the TOPMed imputation reference panel (Taliun et al., 2019) based on hg38 for the phasing and imputation of the SNP array data sets. We mapped the imputed SNPs in hg38 to hg19 using liftover (Hinrichs et al., 2006). The WGS data set of 349 subjects in the MayoRNAseq project was downloaded from Synapse (https://www.synapse.org/#!Synapse:syn10901601). We used the same procedure as in the preparation of the ROSMAP genotype data for the imputation. We imputed the genotype data using the TOPMed imputation reference panel and mapped them to hg19 using liftover. In the allele-specific analysis, we used only high-quality imputed exonic SNPs that had an imputation quality score>0.96 to control phasing errors.

### Analysis of genotype-level and transcript-level *cis*-eQTLs

The genotype-level *cis*-eQTL analysis of the snRNA-seq data was performed for genes whose TSS was within a ±500k bp window of the AD-associated SNPs. Cell-level count matrices of the snRNA-seq data were obtained from the output of STARsolo during the alignment. We included both exonic and intronic reads because pre-mRNA accounts for a large proportion of the snRNA-seq library. We then generated a cell-type-specific subject-level pseudo-bulk count matrix by aggregating cells according to subjects and cell types. We adopted the cell type annotation based on the clustering results in (Habib et al., 2017; Mathys et al., 2019), which are available from Synapse. The pseudo-bulk raw counts were first normalized using the trimmed mean of M-values (TMM) (Robinson and Oshlack, 2010) after removing low-expression genes that had CPM>1 in <3 subjects. Then the overdispersion parameters were estimated using the functions estimateGLMCommonDisp and estimateGLMTagwiseDisp in the edgeR package (Robinson et al., 2010). Finally, the association tests between the gene expression and the genotypes were performed using the glmFit and glmLRT functions with cohorts as the covariate. In a second model to adjust for AD diagnosis, we further added an ordinal variable (between 1 and 5) of the clinical cognitive diagnosis score to the covariates. We also added RNA integrity number (RIN) to the analysis when it is available in the cohort.

For the analysis of transcript-level *cis*-eQTLs in monocytes, we generated a transcript-level count matrix. Specifically, we quantified the transcripts from the fastq files using salmon (Patro et al., 2017) with a pre-built transcriptome index using a partial selective alignment method (http://refgenomes.databio.org/) with the parameters “-l A --validateMappings --rangeFactorizationBins 4”. The downstream analysis followed the same procedure as described above.

### The hierarchical Poisson mixed-effects model for allele-specific eQTL analysis

Testing allelic imbalance is much more powerful than testing individual-level expression data because the allelic information is generally not subject to within-sample technical noises or confounders, which is a severe problem in gene expression study. Here, we propose an HPMM that enjoys the power gain from the allelic imbalance and borrows evidence of the expression of homozygous samples to improve the statistical power further. The HPMM is also straightforward to accommodate covariates and interaction terms. The key idea in the HPMM is to introduce two random-effects terms, which serves two purposes, one is to account for the individual-level overdispersion in the count data, and the other is to convert the two allele counts of each individual to paired samples so that the allelic imbalance within the heterozygous samples has much higher weights than the homozygous individual-level expression.

We first describe the HPMM for analyzing the exonic eQTLs. The model can be applied with a minor modification for the allele-specific eQTL analysis, which will be discussed later. Specifically, each heterozygous subject has two allelic counts corresponding to the two alleles *A* and *a*, and each homozygous subject (*AA* or *aa*) has one count. Denote by *y_ij_* the number of RNA-seq reads overlapping the exonic SNP that had one of the allele patterns *j, j* ∈ {*A, a, AA, aa*} in subject *i, i* ∈ {1,…, *n*}. Thus, for a homozygous subject, we have *j* = *AA or aa* depending on the carried allele, while heterozygous subjects had two data records with *j* = *A* and *a* (Fig. S4a). Denote by *X_ij_* the coding of the allele patterns defined by

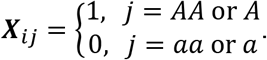

We used the following HPMM to estimate the exonic eQTL effect

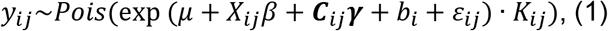

where *μ* is the intercept, *β* is the allele-specific eQTL effect, ***C***_*ij*_ and ***γ*** are the design matrix and effects of covariates, *b_i_* is the individual-level random effects, *ε_ij_* is the allele-level random effects, and *K_ij_* is given by

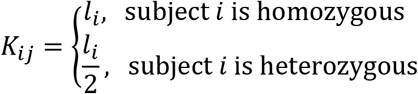

where *l_i_* is the library size or a normalizing factor. That is, *y_ij_* is normalized by half of its total library size of subject *i*. The rationale behind this model is to treat the two records from a heterozygous subject as paired samples by halving its library size and introducing the random effects *b_i_* to capture the individual-level noises, and thus dramatically boosts the power. If there is no homozygous subject, model (1) is equivalent to a binomial model with an overdispersion modeled by *ε_ij_* as shown in (Knowles et al., 2017). The HPMM can be easily fitted using e.g., the glmer function in the lme4 R package (Bates et al., 2014). We performed the exonic eQTL analysis using the glmer function. If the summary statistics show no sign of overdispersion, *ε_ij_* can be omitted to improve the power. To test whether the allelic imbalance was different between two groups (e.g., the AD patients and the control group), we added an interaction term to the model

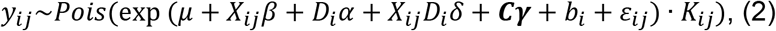

where *D_i_* ∈ {0, 1} is the disease status. Thus, the differential eQTL effects between case-control groups were detected by testing *δ* = 0 in the model (2).

The aseQTL analysis of AD GWAS SNPs adopted the same HPMM framework as that for the exonic eQTLs. The difference is that the allele-specific reads were not directly observed for the GWAS SNPs but were obtained through phased haplotypes as they were not located in an exonic region. We considered haplotypes containing a pair of two SNPs, a GWAS SNP *j* with two alleles *A/a* and an exonic SNP *k* with two alleles *B/b*, where *a* and *b* are the minor alleles. Denote by y¿y the number of reads observed for subject *i* carrying the allele(s) of SNP *j, j* ∈ {*A, a, AA, aa, Aa*}. Note that we split those subjects being heterozygous at both GWAS and exonic loci (i.e., double-heterozygous) into two data records (that is, for subject *i* who is double-heterozygous, we have *j* = {*A, a*}) (Fig. S4b) because allelic reads can be determined for these subjects. For each double-heterozygous subject, we had two data records corresponding to the haplotypes on the two chromosomes (i.e., either (1) *j* = *A, k* = *B* and *j* = *a, k* = *b*, or (2) *j* = *A, k* = *b* and *j* = *a, k* = *B*) and the library size of each data record is halved. Subjects who were not double-heterozygous had only one data record (i.e., *j* = *AA* or *aa* or *Aa*). Note that *j* = *Aa* is used for those who were heterozygous at the GWAS SNP but not at the exonic SNP. Denote by *X_ij_* the coding for the GWAS SNP defined by

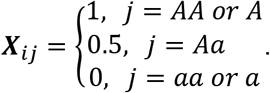

We then fitted model (1) for *X_ij_*, and the effect *β* estimated for *X_ij_* was the aseQTL effect of the AD GWAS SNP.

The interpretation of the allelic analysis of a GWAS SNP is more complicated than that of the exonic aseQTL analysis because the allelic reads are not directly measured for the GWAS SNP. Like GWAS, the LD between the two SNPs complicates the interpretation. Because the double-heterozygous subjects are treated like paired samples in model (1), they have a large impact on the estimate of *β*. Therefore, the haplotype LD in the double-heterozygous subjects is more influential. It should be noted that the haplotype correlation between the GWAS and exonic SNPs in double-heterozygous subjects can be very large even if the two SNPs are not in LD at the population level, particularly when the sample size is small or the MAF is low (Fig. S5). This can also be seen from the following example. Following the previous notations, we consider a situation where there is no LD between the two SNPs, and thus we have that the haplotype relative frequency equals the product of allele frequency, that is, *P_ab_* = *P_a_P_b_*. If the MAF is small (i.e., *P_a_* or *P_b_* is small), then *P_ab_* is close to zero. Thus, when the sample size is small, we may not even observe subjects with the haplotype *ab* in double-heterozygous subjects. The lack of this haplotype means that the only possible combination in double-heterozygous subjects is *j* = *A, k* = *b* and *j* = *a, k* = *B*, which leads to a complete haplotype correlation.

### Quality control in the allele-specific analysis

We performed stringent quality control for sequencing error and mapping bias, which might significantly affect the allele-specific analysis as mentioned in (Castel et al., 2015). As described above, WASP was used in all the analyses, and only those reads with tag vW=1 were retained. In the pileup function, we counted only high-quality reads that were mapped uniquely and had a minimum base quality=10 and minimum MAPQ=254. To minimize potential sequencing and phasing errors, we chose to use the imputed genotypes for the exonic SNPs, but we also double-checked them with the genotypes called from the RNA-seq data. We adopted the RNA-seq called genotypes if the two genotypes were inconsistent. The criterion for the inconsistency was defined by either the ratio between the counts of two alleles >10% when the imputed genotype was homozygous or the ratio <2% when the imputed genotype was heterozygous if >20 reads overlapped the locus in the RNA-seq data. The number of RNA-seq reads at each locus was called using the pileup function in the Rsamtools R package from sorted bam files produced from the fastq files using the Rsubread R package. As the correct phase information is critical for the accuracy of the allele-specific analysis, we examined only those SNPs with an imputation quality score R^2^>0.96.

### Functional annotation and external resource

To compare the eQTL results in CD14+CD16-classical monocytes, we obtained the VCF files containing the summary statistics of significant eQTLs in classical monocytes identified in the DICE. To compared the eQTLs in the brain regions, we obtained the summary statistics of significant eQTLs from GTEx (v8) in three brain tissues, including PFC, hippocampus, and ACC, (The GTEx Consortium, 2020), and from a meta-analysis of TC and PFC conducted in (Sieberts et al., 2020). The summary statistics of sQTLs in the PFC are obtained from a supplementary table of (Raj et al., 2018). The functional annotation of the exonic variants in the aseQTL analysis was performed using the VariantAnnotation R package (Obenchain et al., 2014), in which we annotated the location of the exonic variants using the database *TxDb.Hsapiens.UCSC.hg38.knownGene*, the amino acid coding change (non-synonymous, synonymous, frameshift, and nonsense), and prediction of the impact of non-synonymous variants using PolyPhen (Polymorphism Phenotyping).

## Acknowledgements

This manuscript was prepared using limited access datasets obtained through Synapse and dbGaP (accession numbers: phs00424.v5.p1 (GTEx), phs001703.v3.p1 (DICE)).

This research was supported by Grants from the National Institutes of Health R01 AG061853, R01 AG065477, and R01 AG070488 to A.M.K.

The funders had no role in study design, data collection, and analysis, decision to publish, or manuscript preparation. The content is solely the responsibility of the authors and does not necessarily represent the official views of the National Institutes of Health.

We are grateful to the participants in the Religious Order Study, the Rush Memory and Aging Project. Data can be requested at www.radc.rush.edu.

The results published here are in whole or in part based on data obtained from the AD Knowledge Portal (https://adknowledgeportal.org). Study data were provided by the following sources: The Mayo Clinic Alzheimers Disease Genetic Studies, led by Dr. Nilufer Taner and Dr. Steven G. Younkin, Mayo Clinic, Jacksonville, FL using samples from the Mayo Clinic Study of Aging, the Mayo Clinic Alzheimers Disease Research Center, and the Mayo Clinic Brain Bank. Data collection was supported through funding by NIA grants P50 AG016574, R01 AG032990, U01 AG046139, R01 AG018023, U01 AG006576, U01 AG006786, R01 AG025711, R01 AG017216, R01 AG003949, NINDS grant R01 NS080820, CurePSP Foundation, and support from Mayo Foundation. Study data includes samples collected through the Sun Health Research Institute Brain and Body Donation Program of Sun City, Arizona. The Brain and Body Donation Program is supported by the National Institute of Neurological Disorders and Stroke (U24 NS072026 National Brain and Tissue Resource for Parkinsons Disease and Related Disorders), the National Institute on Aging (P30 AG19610 Arizona Alzheimers Disease Core Center), the Arizona Department of Health Services (contract 211002, Arizona Alzheimers Research Center), the Arizona Biomedical Research Commission (contracts 4001, 0011, 05-901 and 1001 to the Arizona Parkinson’s Disease Consortium) and the Michael J. Fox Foundation for Parkinsons Research.

## Author contributions

LH conceived the study. LH developed the statistical model, preprocessed the data, and performed the analysis. YL contributed to data preprocessing. AK contributed to acquiring the data, and discussion of the final results. All authors contributed to the writing of the manuscript.

## Supplementary Tables

Table S1. Summary statistics of the analysis of aseQTLs for the AD-associated SNPs in the four brain regions using the bulk RNA-seq data. AD SNP: the rs ID of the AD-associated SNP. AD SNP Info: the detailed information of the AD-associated SNP including chromosome, genomic location (hg19), reference allele, and effect allele. Exonic SNP Info: the detailed information of the exonic variant used for measuring the ASE. Mean expression count: the mean raw count of the expression measured at the exonic variant. Beta (P) of AD SNP: the logFC (p-value) of the effect of the AD-associated SNP on the expression. Beta (P) of exonic SNP: the logFC (p-value) of the effect of the exonic variant on the expression. Sample size: Actual sample size in testing this association. Gene name: the gene where the exonic SNP is located. Amino acid modification, Exonic region, Prediction for nonsynonymous: Functional annotation of the exonic SNP. Padj: FDR adjusted p-value.

Table S2. Summary statistics of the interaction analysis between age and genotype for the significant AD-associated SNPs detected in the four brain regions using the bulk RNA-seq data. Exonic SNP pos: the genomic position (hg19) of the exonic variant used for measuring the ASE. Mean count: the mean raw count of the expression measured at the exonic variant. P of interaction: the p-value of the interaction term between age and genotype.

Table S3. Summary statistics of the analysis of aseQTLs for the AD-associated SNPs in the six neural cell types using the snRNA-seq data. AD SNP: the rs ID of the AD-associated SNP. AD SNP Info: the detailed information of the AD-associated SNP including chromosome, genomic location (hg19), reference allele, and effect allele. Exonic SNP Info: the detailed information of the exonic variant used for measuring the ASE. Mean expression count: the mean raw count of the expression measured at the exonic variant. Beta (P) of AD SNP: the logFC (p-value) of the effect of the AD-associated SNP on the expression. Beta (P) of exonic SNP: the logFC (p-value) of the effect of the exonic variant on the expression. Gene name: the gene where the exonic SNP is located. Amino acid modification, Exonic region, Prediction for nonsynonymous: Functional annotation of the exonic SNP. Padj: FDR adjusted p-value.

Table S4. Summary statistics of the analysis of subject-level *cis*-eQTLs for the AD-associated SNPs in the six neural cell types using pseudo-bulk samples by aggregating cells in the snRNA-seq data. logCPM: logarithm of the count per million of the gene expression. Adjusted P: FDR adjusted p-value. SNP: the SNP ID, the effect allele (first), and the reference allele (second).

Table S5. Summary statistics of the analysis of aseQTLs for the AD-associated SNPs in microglia using the combined snRNA-seq and cell-sorting bulk RNA-seq data. AD SNP: the rs ID of the AD-associated SNP. AD SNP Info: the detailed information of the AD-associated SNP including chromosome, genomic location (hg19), reference allele, and effect allele. Exonic SNP Info: the detailed information of the exonic variant used for measuring the ASE. Mean expression count: the mean raw count of the expression measured at the exonic variant. Beta (P) of AD SNP: the logFC (p-value) of the effect of the AD-associated SNP on the expression. Beta (P) of exonic SNP: the logFC (p-value) of the effect of the exonic variant on the expression. Sample size: Actual sample size in testing this association. Gene name: the gene where the exonic SNP is located. Amino acid modification, Exonic region, Prediction for nonsynonymous: Functional annotation of the exonic SNP. Padj: FDR adjusted p-value.

Table S6. Summary statistics of the analyses of subject-level *cis*-eQTLs for the AD-associated SNPs in microglia using pseudo-bulk samples by aggregating cells in the combined snRNA-seq and cell-sorting bulk RNA-seq data. The first analysis is adjusted for batch, and the second analysis is adjusted for both batch and the diagnosis of AD. logCPM: logarithm of the count per million of the gene expression. Adjusted P: FDR adjusted p-value. SNP: the SNP ID, the effect allele (first), and the reference allele (second).

Table S7. Summary statistics of the analysis of aseQTLs for the AD-associated SNPs in monocytes using the cell-sorting bulk RNA-seq data. AD SNP: the rs ID of the AD-associated SNP. AD SNP Info: the detailed information of the AD-associated SNP including chromosome, genomic location (hg19), reference allele, and effect allele. Exonic SNP Info: the detailed information of the exonic variant used for measuring the ASE. Mean expression count: the mean raw count of the expression measured at the exonic variant. Beta (P) of AD SNP: the logFC (p-value) of the effect of the AD-associated SNP on the expression. Beta (P) of exonic SNP: the logFC (p-value) of the effect of the exonic variant on the expression. Sample size: Actual sample size in testing this association. Gene name: the gene where the exonic SNP is located. Amino acid modification, Exonic region, Prediction for nonsynonymous: Functional annotation of the exonic SNP. Padj: FDR adjusted p-value.

Table S8. Summary statistics of the eQTL analysis of rs3865444 with the transcript-level expression of *CD33*.

## Supplementary Figures

Figure S1. Allelic and genotype-level expression plots of the eight significant associations that were identified in the aseQTL analysis in monocytes and are not shown in Fig. 6. The boxplots summarize the subject-level expression and the ASE of the double-heterozygous samples (yellow points) are shown in the scatter plot below. The dashed line is the smooth curve fitted using linear regression.

Figure S2. Three significant associations identified in the aseQTL analysis in microglia between (a) rs59735493 and *ITGAM*, (b) rs2081545 and *MS4A7*, and (c) rs12590654 and *ATXN3*. ASE_NA indicates those samples that ASE is not available because they are heterozygous but not double-heterozygous. The pair of the ASE of a double-heterozygous sample is connected by a dashed line. The CPM of the ASE of the double-heterozygous sample was normalized by half of its total library size.

Figure S3. Significant associations between (a) rs1859788 and the ASE of *TRIM4*, and (b) rs3740688 and the ASE of *MTCH2* in the excitatory neurons. The boxplots summarize the subject-level expression and the ASE of the double-heterozygous samples (yellow points) are shown in the scatter plot. The dashed line is the smooth curve fitted using linear regression.

Figure S4. The coding strategy used in the proposed HPMM for the aseQTL analysis of the association (a) between an exonic SNP and its ASE and (b) between a GWAS SNP and the ASE measured at the exonic SNP.

Figure S5. The difference between the genotype correlation among the whole sample and the haplotype correlation among double-heterozygous subjects observed in the 367 significant associations (FDR p<0.05) identified in the analysis of aseQTLs in the four brain regions. X axis: the product of the MAF of the AD-associated SNP and the exonic SNP. The color of the points indicates whether the genotype correlation among the whole sample is positive or negative.

