## Supplementary figures and images for "Allele-specific analysis reveals exon- and cell-type-specific regulatory effects of Alzheimer’s disease-associated genetic variants"

### Figure S1

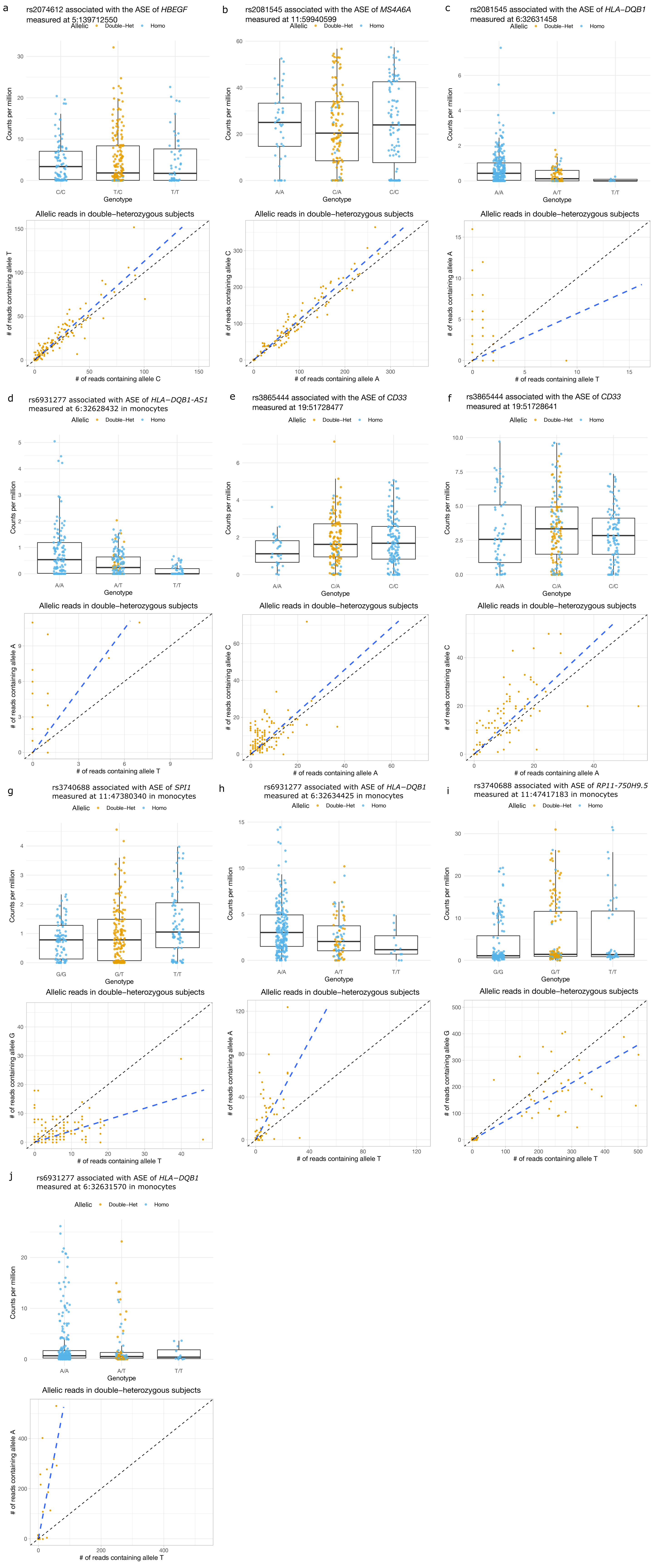

### Figure S2

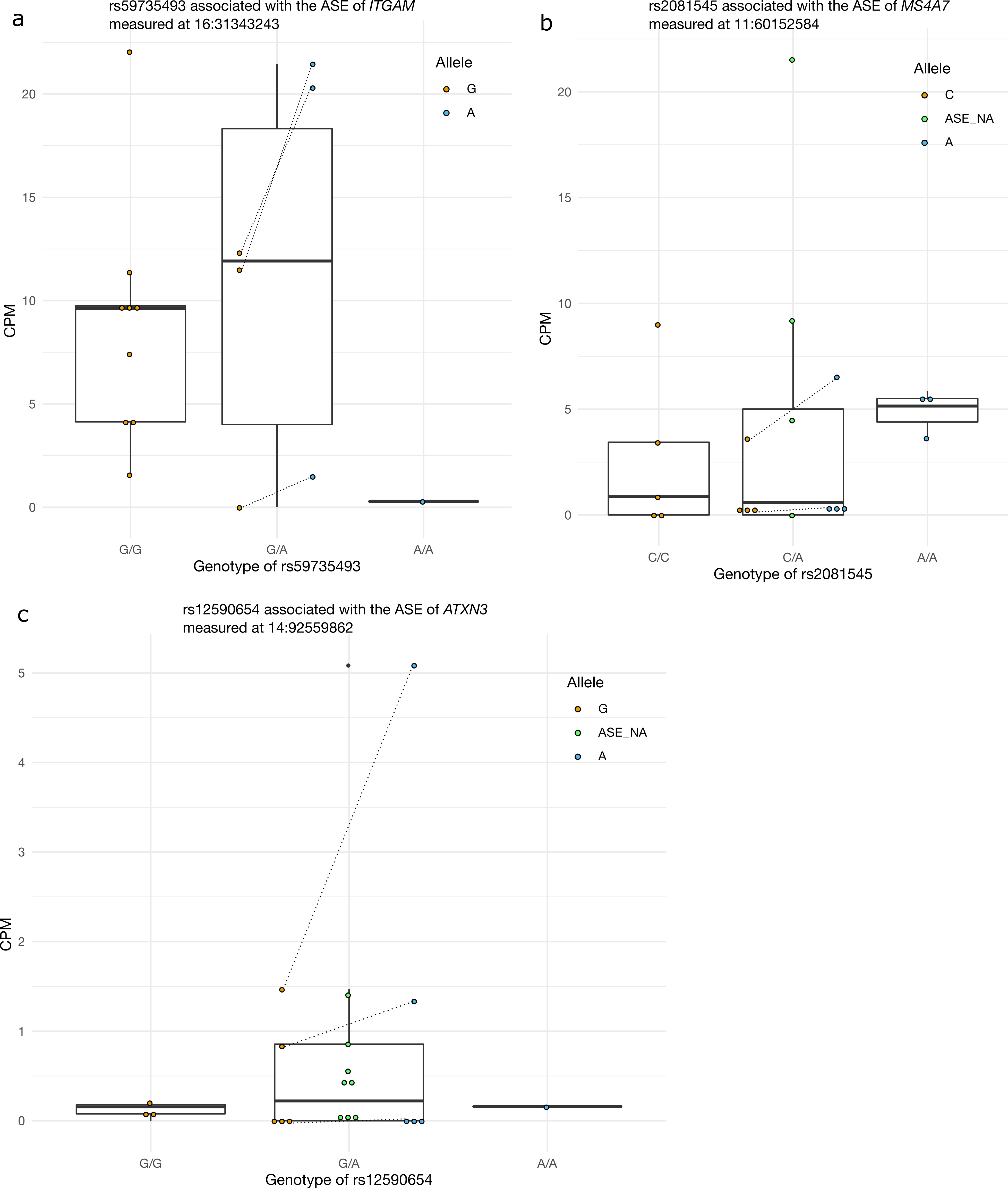

### Figure S3

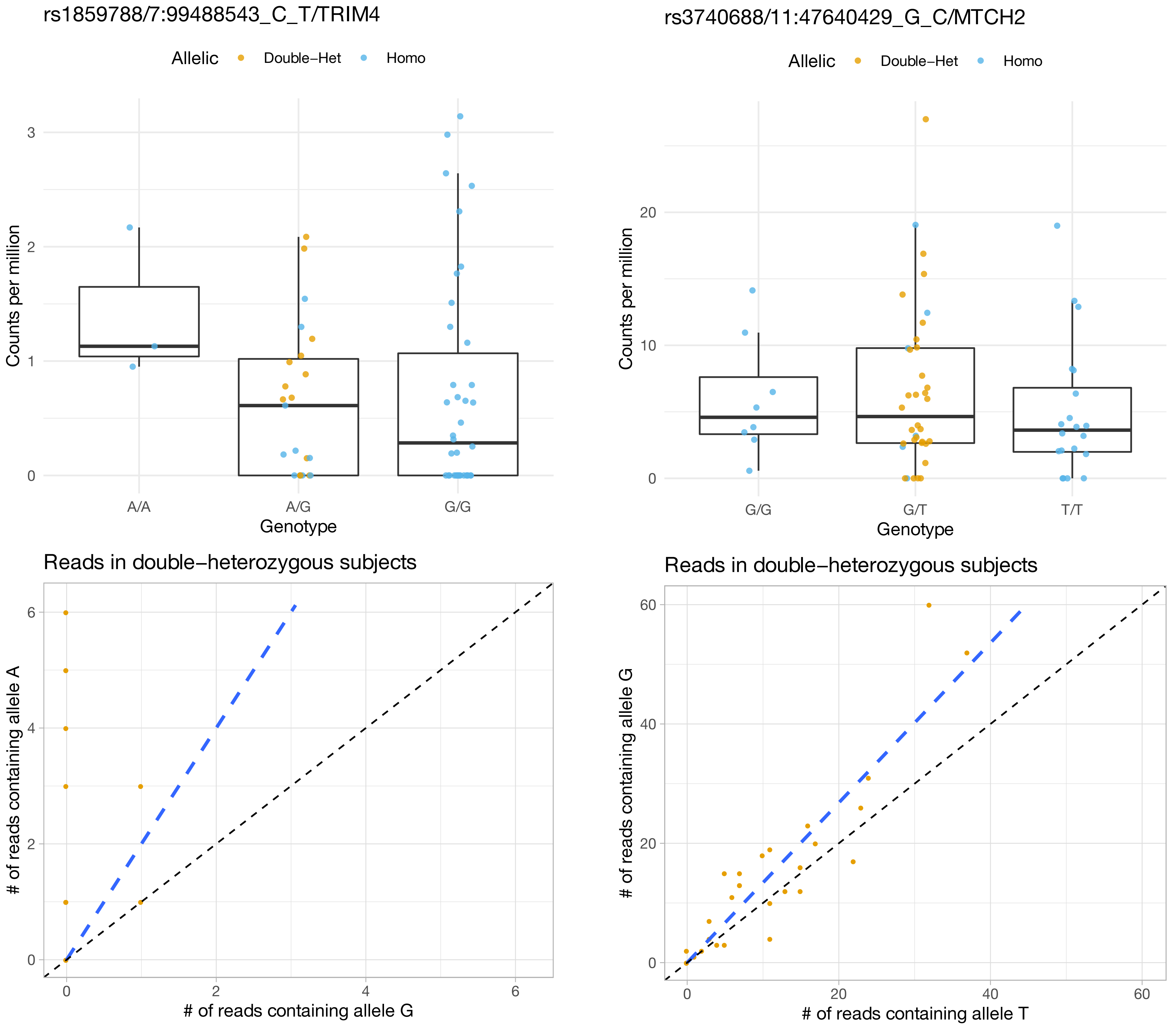

### Figure S5

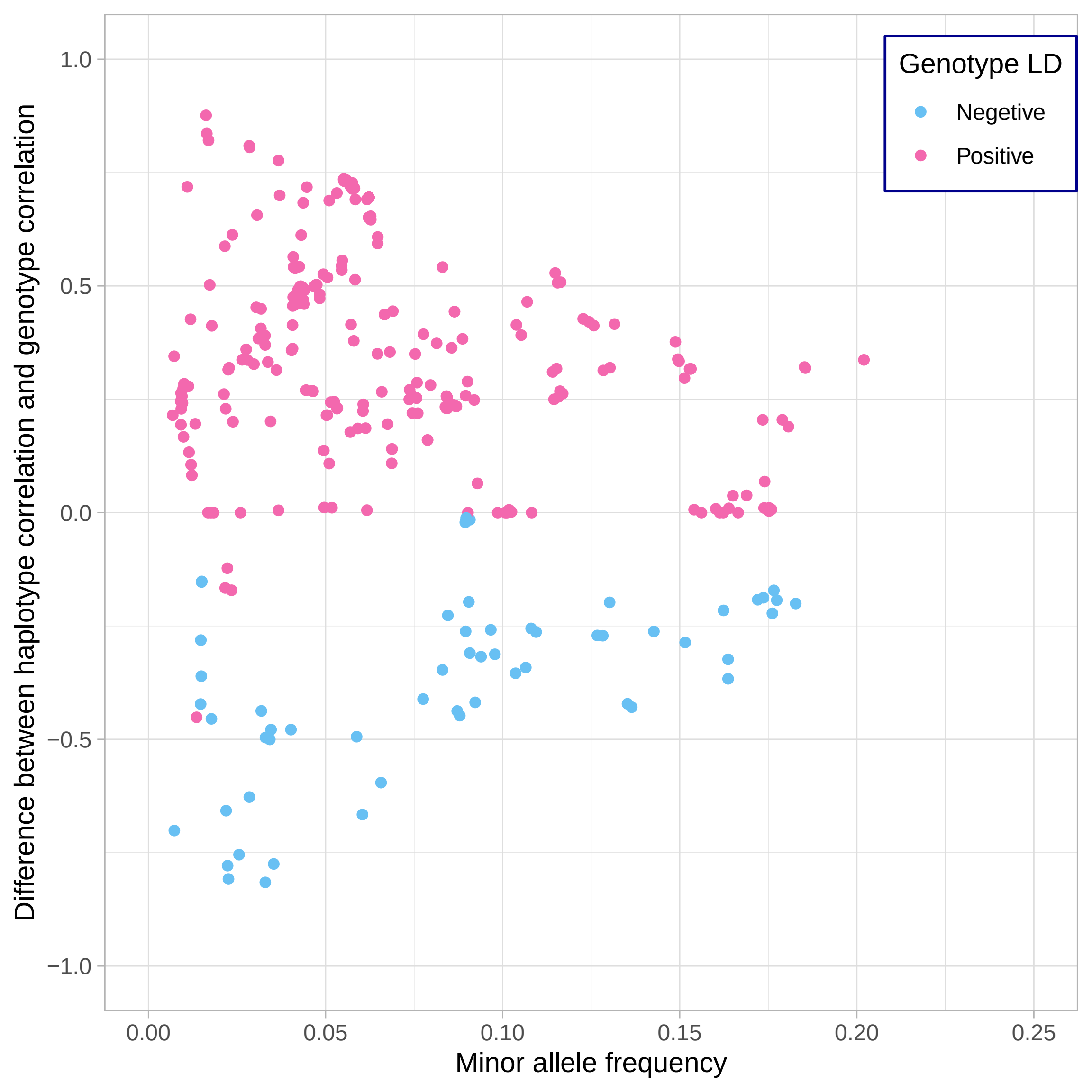
