## Supplementary material for "Allele-specific analysis reveals exon- and cell-type-specific regulatory effects of Alzheimer’s disease-associated genetic variants": Figure S4

a Coding strategy for testing allelic imbalance for an exonic SNP of which the ASE is measured

Genotype-level data set

| Subject ID | Genotype | Dosage | Count | Library |
|---|---|---|---|---|
| 1 | A/A | 0 | 5 | 1000 |
| 2 | A/a | 1 | 14 | 1000 |
| 3 | a/a | 2 | 10 | 1000 |

Allelic expression available for the heterozygous subject

Allele-specific data set

| Subject ID | Allele | Allele coding | Count | Library |
|---|---|---|---|---|
| 1 | A | 0 | 5 | 1000 |
| 2 | A | 0 | 5 | 500 |
| 2 | a | 1 | 9 | 500 |
| 3 | a | 1 | 10 | 1000 |

b Coding strategy for testing allelic imbalance for a GWAS SNP through the haplotype with an exonic SNP

| Subject ID | GWAS SNP | Exonic SNP | Count | Library |
|---|---|---|---|---|
| 1 | A/A | B/b | 5 | 1000 |
| 2 | A/a | B/b | 12 | 1000 |
| 3 | A/a | B/B | 9 | 1000 |
| 4 | a/a | B/B | 10 | 1000 |

Allelic expression available for the double-heterozygous subject

| Subject ID | GWAS Allele | GWAS coding | Count | Library |
|---|---|---|---|---|
| 1 | A | 0 | 5 | 1000 |
| 2 | A | 0 | 5 | 500 |
| 2 | a | 1 | 7 | 500 |
| 3 | A/a | 0.5 | 9 | 1000 |
| 4 | a | 1 | 10 | 1000 |
